# Single-nucleus and spatial transcriptomic profiling of human temporal cortex and white matter reveals novel associations with AD pathology

**DOI:** 10.1101/2024.04.23.590816

**Authors:** Pallavi Gaur, Julien Bryois, Daniela Calini, Lynette Foo, Jeroen J M Hoozemans, Dheeraj Malhotra, Vilas Menon

## Abstract

Alzheimer’s disease (AD) is a neurodegenerative disorder with complex pathological manifestations and is the leading cause of cognitive decline and dementia in elderly individuals. A major goal in AD research is to identify new therapeutic pathways by studying the molecular and cellular changes in the disease, either downstream or upstream of the pathological hallmarks. In this study, we present a comprehensive investigation of cellular heterogeneity from the temporal cortex region of 40 individuals, comprising healthy donors and individuals with differing tau and amyloid burden. Using single-nucleus transcriptome analysis of 430,271 nuclei from both gray and white matter of these individuals, we identified cell type-specific subclusters in both neuronal and glial cell types with varying degrees of association with AD pathology. In particular, these associations are present in layer specific glutamatergic (excitatory) neuronal types, along with GABAergic (inhibitory) neurons and glial subtypes. These associations were observed in early as well as late pathological progression. We extended this analysis by performing multiplexed in situ hybridization using the CARTANA platform, capturing 155 genes in 13 individuals with varying levels of tau pathology. By modeling the spatial distribution of these genes and their associations with the pathology, we not only replicated key findings from our snRNA data analysis, but also identified a set of cell type-specific genes that show selective enrichment or depletion near pathological inclusions. Together, our findings allow us to prioritize specific cell types and pathways for targeted interventions at various stages of pathological progression in AD.

## Main

The two pathological hallmarks of Alzheimer’s Disease (AD) - amyloid-beta plaques and neurofibrillary tangles - show accumulation and spread that is associated with the severity of symptoms ^1,2,3^. Although the complex pathophysiology of AD remains poorly understood, recent transcriptomic and epigenomic analyses have revealed that post-mortem human AD brain tissue exhibits downregulation of genes associated with neuronal function^4,5^ and upregulation of the genes involved in the innate immune response^6,7^. However, not many studies have looked systematically at the spatial distribution of gene expression and its relationship to neuropathology^8^. Thus, our overall understanding of cell type heterogeneity and compositional changes during pathological accumulation is still under-explored, hindering our ability to understand the biological processes underlying AD^9^.

While the pathogenesis of Alzheimer’s disease (AD) has been extensively studied, the predominant focus has traditionally been on gray matter alterations, overlooking the essential role that white matter plays in neurological health. Recognizing the involvement of the temporal cortex in neurodegenerative diseases is pivotal for developing precise interventions and therapies. Here we investigate the cellular heterogeneity in the white and gray matter of the temporal cortex (TC) region of 40 individuals taken from Netherlands brain bank (NBB), comprising healthy donors and individuals at varying Braak stages. We obtained 430,271 single-nucleus RNA profiles from white and gray matter of the TC and deployed a comprehensive analysis approach to identify associations with varying degrees of AD pathology. To further expand our analysis of transcriptomic signatures within the TC, we incorporated data from five previously published studies, including 888,784 total nuclei (Table1). This comprehensive analysis with different brain regions such as the entorhinal cortex^10,11^, prefrontal cortex^12,13,5^, and superior frontal gyrus^10^ allowed us to replicate a subset of the cell type associations. Finally, we explored the spatial distribution of a subset of key cell type-specific genes in 13 individuals through the CARTANA targeted in-situ sequencing (ISS) platform^14^ (Fig.1a). Starting with a panel of 155 genes, we investigated spatial enrichment or depletion with Braak stage (validating our snRNA-seq results), as well as enrichment or depletion near specific pathological inclusions identified on the same tissue sections.

**Fig.1:**
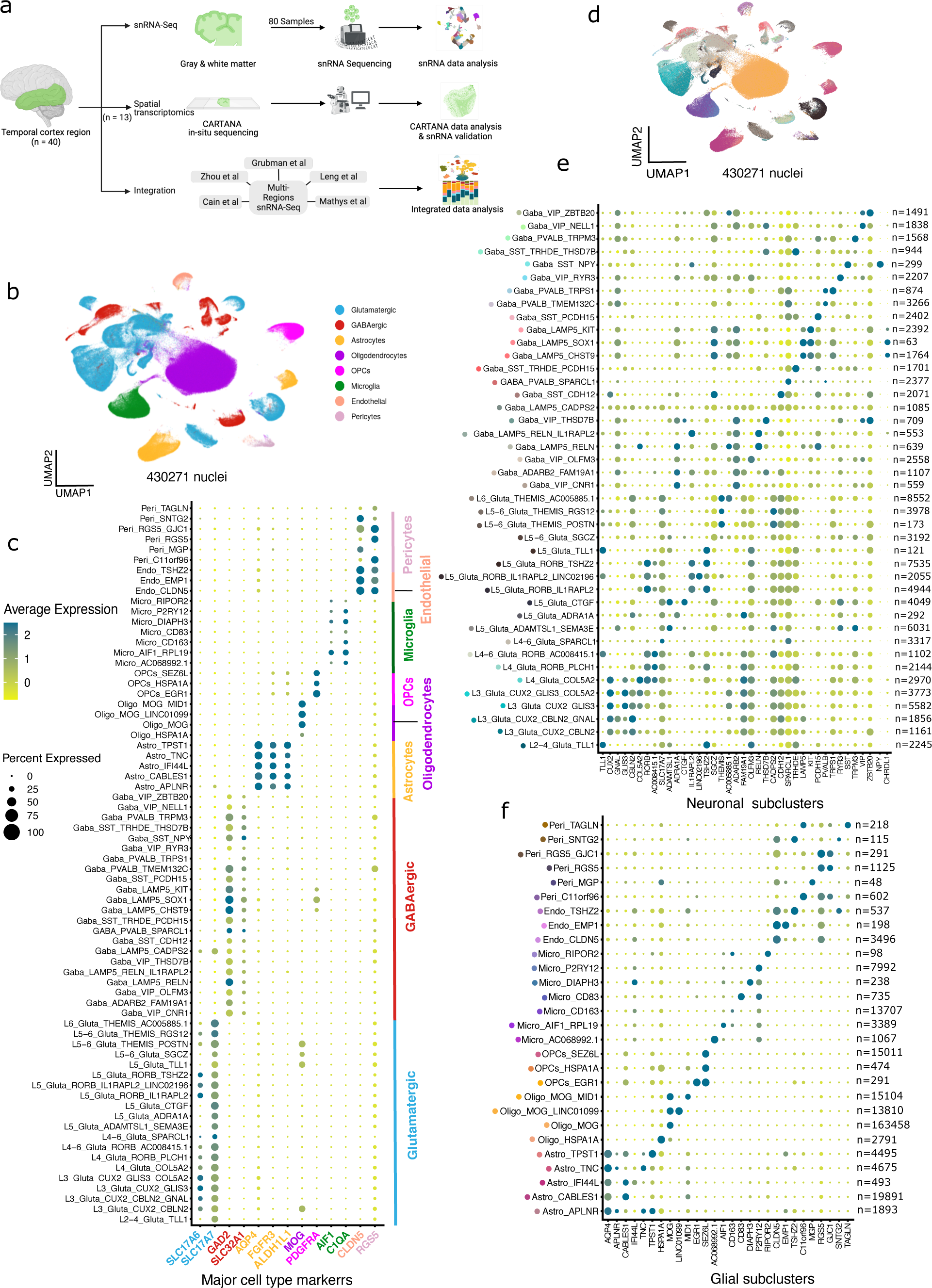
Experimental scheme and molecular map of the human TC in 40 individuals (controls and AD) **(a)** Overview of the experimental scheme **(b)** UMAP embedding of 430,271 single-nucleus RNA profiles from the TC brain region of 40 individuals; colored by cell type **(c)** Dot plot of the canonical markers distinguishing 8 major cell types with different levels of expression (color) and percentage of expressing nuclei (dot size) across 430,271 TC nuclei **(d)** UMAP embedding of 430,271 single-nucleus RNA profiles from the TC brain region colored by different subclusters **(e)** Dot plot of the DE markers distinguishing different neuronal cell subtypes with different levels of expression and percentage of expressing nuclei across different subclusters with numbers of nuclei (n) shown in right **(f)** Dot plot of the DE markers distinguishing different glial cell subtypes with different levels of expression and percentage of expressing nuclei across different subclusters with numbers of nuclei (n) shown in right.

**Table 1.**

|  | **Brain region** | **Total No. of Samples** | **Total No. of nuclei** |
| --- | --- | --- | --- |
| **Roche (TC)** | Temporal Cortex/ White Matter | 80 (40*2) | 430271 |
| **Cain et al** | Prefrontal Cortex/White matter | 48 (24*2) | 226951 |
| **Zhou et al** | Prefrontal Cortex | 32 | 89839 |
| **Mathys et al** | Prefrontal Cortex | 48 | 43281 |
| **Leng et al** | Entorhinal Cortex/Superior Frontal Gyrus | 20(10*2) | 81098 |
| **Grubman et al** | Entorhinal Cortex | 12 | 17344 |

## Results

### Transcriptional states of CNS cell types in the temporal cortex gray and white matter of the aged human brain

We profiled tissues from the temporal cortex region to investigate the association of pathology with different sub cell types proportions. To achieve this, first we profiled the transcriptome of 80 post-mortem human brain samples (40 individuals; gray and white matter) encompassing controls and individuals with varying degrees of AD pathology (Extended Data Fig.1a, b). We retained 430,271 nuclei for AD trait association analysis after quality control and cluster identification. Nuclei segregated into eight major cell types: glutamatergic neurons, GABAergic neurons, microglia, astrocytes, oligodendrocytes, oligodendrocyte precursor cells (OPCs), endothelial cells, and pericytes (Fig.1b). Nuclei in these broad cell types showed specific expression of canonical marker genes such as *SLC17A6* and *SLC17A7* for glutamatergic neurons, *GAD1* and *SLC32A1* for GABAergic neurons, *AQP4, FGFR3* and *ALDH1L1* for astrocytes, MOG for oligodendrocytes, *PDGFRA* for OPCs, *AIF1* and *C1QA* for microglia, *CLDN5* for endothelial cells, and *RGS5* for pericytes (Fig.1c). Each major cell type was then subclustered independently, and subclusters were named based on a combination of top differentially expressed markers (Methods, Fig.1d, e, f). We identified 20 subpopulations of glutamatergic neurons, 22 subpopulations of GABAergic neurons, 5 of astrocytes, 4 of oligodendrocytes, 3 of OPCs, 7 of microglia, 3 of endothelial cells and 6 of pericytes (Supplementary Table1).

### Neuronal and glial subpopulations with lower prevalence in late Braak stages

Our key analysis was to associate differences in subpopulation composition within each major cell type with specific AD phenotypes. We selected neurofibrillary tangle (NFT) spread as the AD phenotype of interest because of its stereotyped progression and its strong association with cognitive decline^3,12^. In particular, we used the Braak staging paradigm, which assigns a stage (from 1 to 6) based on the overall spread of tau pathology in an individual, with early stages (Braak 1/2) indicating NFT accumulation primarily in the entorhinal cortex and hippocampus, and later stages (Braak 3/4/5/6) describing individuals with tau pathology in both early and late-affected (cortical) regions. Individuals at Braak stage 3 and beyond tend to show NFT accumulation in temporal cortex, a fact we verified with staining for tau in 13 of our donors (Supplementary Table1) as part of our CARTANA experiments. With Braak stage as our primary pathology variable, we then examined cell subpopulations that showed coordinated changes with this variable.

In the gray matter (GM), we identified multiple subsets of neurons showing lower proportions in individuals at late Braak stages (Fig.2a, b). Out of 5 total *RORB+* glutamatergic subpopulations, two were found to be less prevalent in later Braak stages compared to earlier stages; interestingly, both of these *RORB+* subpopulations express *IL1RAPL2*. This aligns with recent findings where *RORB* has been recognized as an indicative marker of selectively vulnerable excitatory neurons in the entorhinal cortex^10^, although we find that not all *RORB+* subpopulations are equally vulnerable at late Braak stages. We also observed a glutamatergic subset with high levels of *SPARCL1* was lower in the GM in donors at advanced Braak stages^15^. Other subpopulations with lower proportions at later Braak stages included those marked by *COL5A2* (putatively in Layer 4) and *GLIS3* (putatively in Layer 3). Prior microarray-based studies have found *COL5A2* to be downregulated in AD^16,17^, and *GLIS3* has been implicated in a genome-wide significant association with cerebrospinal fluid tau and phospho-tau levels in the context of a specific genetic variant^18^. Finally, we also observed lower proportions of *GNAL+* and *TLL1+* glutamatergic subpopulations at Braak stages 5/6. The glutamatergic subpopulation expressing *GNAL* has been reported to have alternative splicing association with AD^19^ and *TLL1* has been reported as one of the five genes (*Tshz2, Gm12695, St3gal1, Isx and Tll1*) implicated in Aβ processing, and is affected in Tg-5xFAD mice treated with REMFS^20^.

**Fig.2:**
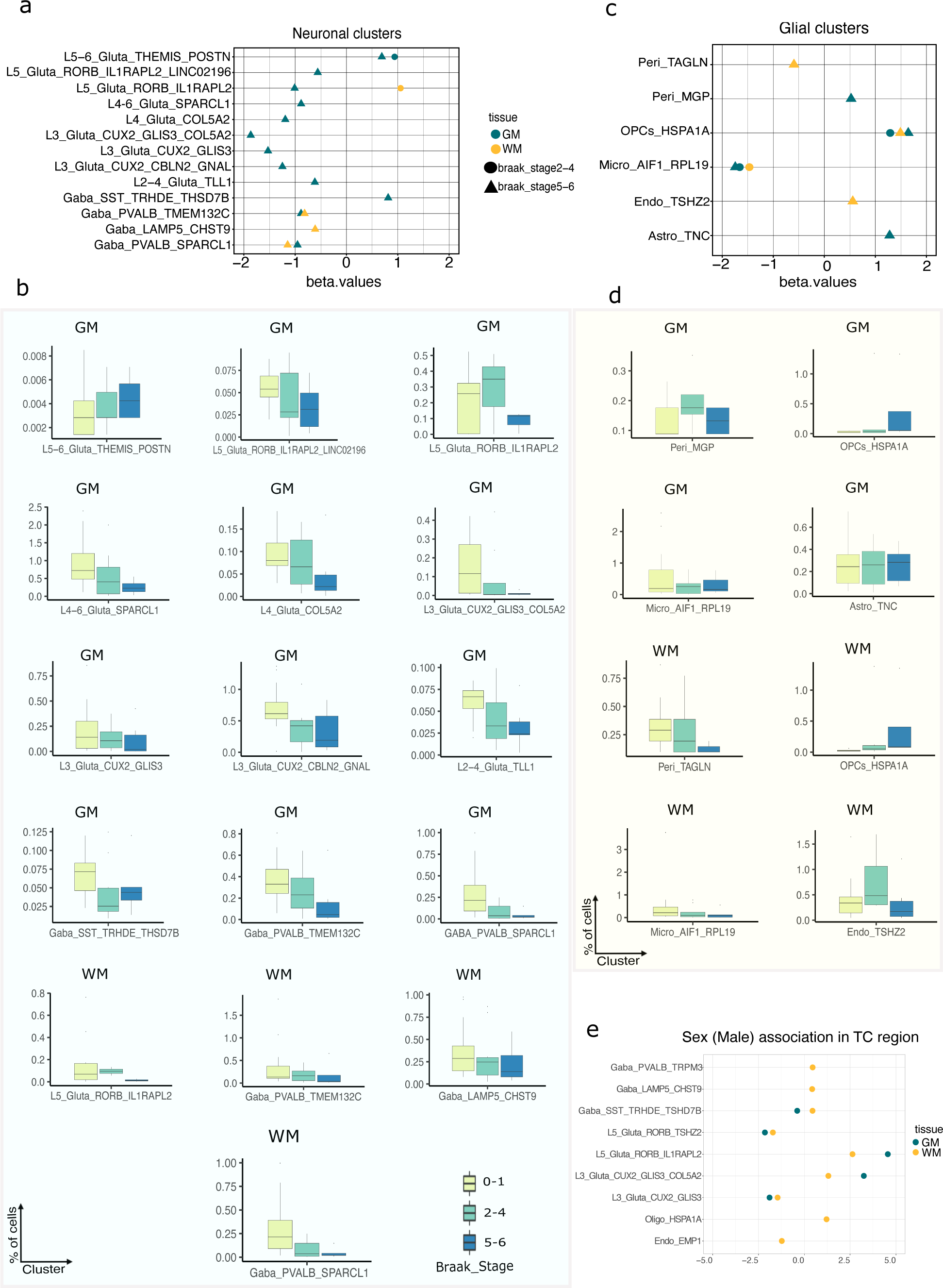
AD trait association analysis of different cellular subtypes of TC region. **(a)** Differential abundance of statistically significant neuronal subpopulations with respect to AD pathology indicated by varying Braak stages in GM and WM. The x-axis represents the estimated log-ratio differences in relative abundance between different Braak stages and the y-axis represents different neuronal subpopulations. **(b)** Box plots of all neuronal subpopulations shown in panel a, representing the distribution of nuclei in each subpopulation with respect to their concerned major cell type. **(c)** Differential abundance of statistically significant glial subpopulations with respect to AD pathology indicated by varying Braak stages. The x and y axis representations are the same as the panel ‘a’. (**d)** Box plots of all glial subpopulations shown in panel ‘c’. **(e)** Differential abundance of statistically significant subpopulations with respect to sex (male).

With respect to GABAergic neurons in GM, we found multiple subgroups with lower proportions at late Braak stages. We observed that parvalbumin (*PVALB*) neurons have lower proportions in GM in individuals at late Braak stages (Fig.2a). *PVALB+* neurons have been reported to be selectively depleted in the frontal cortex of Alzheimer’s disease mouse models^21^. Interestingly, *SPARCL1+* GABAergic neurons also showed depleted proportions in GM. Notably, *SPARCL1* has been reported to have altered levels in the cerebrospinal fluid (CSF) of AD patients and it is conceivable that alterations in the regulation of *SPARCL1* may play a significant role in the development of AD^22,23^. On the glial side, we found associations between microglial sub-signatures with later Braak stages. Specifically, *RPL19+* microglia were significantly lower in intermediate and late Braak stages in GM, possibly reflecting the loss of a protective type of microglia (Fig.2c,d).

### Specific neuronal subtypes show higher proportions in GM at late Braak stages

Our analysis above highlighted multiple neuronal groups with lower proportions at late Braak stages. However, since these proportions are measured as a fraction of the total glutamatergic or GABAergic neurons, they are offset by concomitantly higher proportions of other neuronal subtypes; these latter groups are putatively less vulnerable to disease pathology and are interesting for further study. For example, we found that a few glutamatergic neuron populations from the deep layers of the cortex (L5−6) in GM, marked by *THEMIS* and *POSTN*, are overrepresented at later Braak stages, which suggests that this subtype could be more resistant to increasing tau burden. In addition, a unique *SST/THSD7B+* GABAergic subpopulation marked by Thyrotropin Releasing Hormone Degrading Enzyme (*TRHDE*) was also enriched in Braak stage 5/6 (Fig.2a,b). Among glial cells, OPCs specific to heat shock protein were enriched during the AD pathology progression in GM. A similar pattern was observed in *MGP+* pericytes and *TNC+* astrocytes, whose proportions were higher in individuals at later Braak stages.

### A subset of cell clusters shows changes in temporal cortex associated white matter

Although the vast majority of cells in the white matter are oligodendrocytes, we found changes in glial subpopulations, and even some neuronal populations, associated with Braak stage in TC-associated white matter (WM). Endothelial cells marked by *TSHZ2* were present at higher proportions in donors at advanced Braak stages (Fig.2c), as were OPCs with high expression of heat-shock proteins; this latter finding resembles the same finding in GM. Finally, *RPL19+* microglia and *TAGLN+* pericytes were lower in individuals at intermediate and advanced Braak stages, respectively; indeed, *RPL19+* microglia were one of the cell signatures showing the same trend in WM and GM.

Despite the sparsity of neuronal nuclei in the WM, we observed statistically significant differences in neuronal subgroups in WM at different Braak stages. Similar to our findings in GM, *PVALB+/TMEM132C+* and *PVALB+/SPARCL1+* subpopulations were lower in WM in donors at advanced Braak stages (Fig.2a,b). By contrast, *RORB+/IL1RAPL2+* glutamatergic subpopulations showed a different pattern in WM, exhibiting higher proportions at intermediate Braak stages 2/4 (Fig.2a,b). Finally, GABAergic neurons expressing *LAMP5 -* a gene previously implicated in dysfunction in Alzheimer’s disease^24^ - and *CHST9*, a drug metabolism-related gene in AD^25^, showed lower prevalence in WM in late Braak stages (5-6) (Fig.2a,b). Overall, the changes in WM appear to be less distinct than those found in GM and are mostly restricted to rarer cell populations.

### Consistency of subcluster associations with AD pathology traits across multiple brain regions and studies

In order to assess the generalizability of our results, we integrated our snRNA-Seq data with other publicly available datasets spanning various brain regions. We incorporated entorhinal cortex^10,11^, prefrontal cortex^5,12,13^, and superior frontal gyrus^10^ and deep white matter from prefrontal cortex^12^ to examine the single nuc RNA-Seq profiles (Extended Data Fig.2a,b, Table1). After extensive quality control and filtering of 959,237 nuclei (Methods), we retained a total of 888,784 nuclei for trait association analysis (Supplemental Table1). We then investigated which AD trait associations in the TC were found across multiple brain regions (Supplemental Table1). We found that *THEMIS+/POSTN+* deep layer glutamatergic neurons were consistently overrepresented in late Braak stages, suggesting these deep layer neurons may be generally resistant to AD pathology across the cortex (Extended Data Fig.3c). Conversely, *PVALB+/TMEM132C+* GABAergic neurons were lower at advanced Braak stages in all brain regions we examined (Extended Data Fig.3c). On the glial side, *TAGLN+* pericytes and *RPL19+* microglia were lower in Braak stages 5/6 in multiple cortical regions, whereas *HSPA1A+* OPCs were enriched in donors at these Braak stages (Extended Data Fig.3d). We also investigated the AD trait associations exclusively in pre-frontal cortex (PFC) region and observed the same consistent overrepresentation of *THEMIS+/POSTN+* deep layer glutamatergic neurons in late Braak stages (Extended Data Fig.3e). Along with this, *TAGLN+* pericytes were also lower in advanced Braak stages (Extended Data Fig.3f) in PFC.

### Differential expression analysis identifies cell type and tissue specific changes in late stage AD pathology in the TC

Whereas cluster proportion analyses can identify differential vulnerability and resistance, disease-associated cellular signatures may be obscured when cluster boundaries are not discrete. To identify additional signatures potentially missed by cluster-based analyses, we then looked for pathological changes in gene expression levels in each broad cell type using a pseudobulk approach (Methods). As our experimental design included two samples per individual (GM matter from the TC + associated WM), we used a mixed effect negative binomial model to test for changes in gene expression related to AD pathology (Methods). We found few expression changes in the early stages of AD (Braak 0-1 vs Braak 2-4), with only 17 genes differentially expressed (FDR-adjusted p-value <0.05), and all of these were in GABAergic neurons, oligodendrocytes, and OPCs (Fig.3a). As our statistical model included an interaction term between Braak stage and tissue, we also tested whether any genes were differentially affected by early AD pathology in gray vs white matter for each broad cell type. We found only a few genes that were affected at early Braak stage differentially between GM and WM (10 genes, FDR-adjusted p-value <0.05).

**Fig.3:**
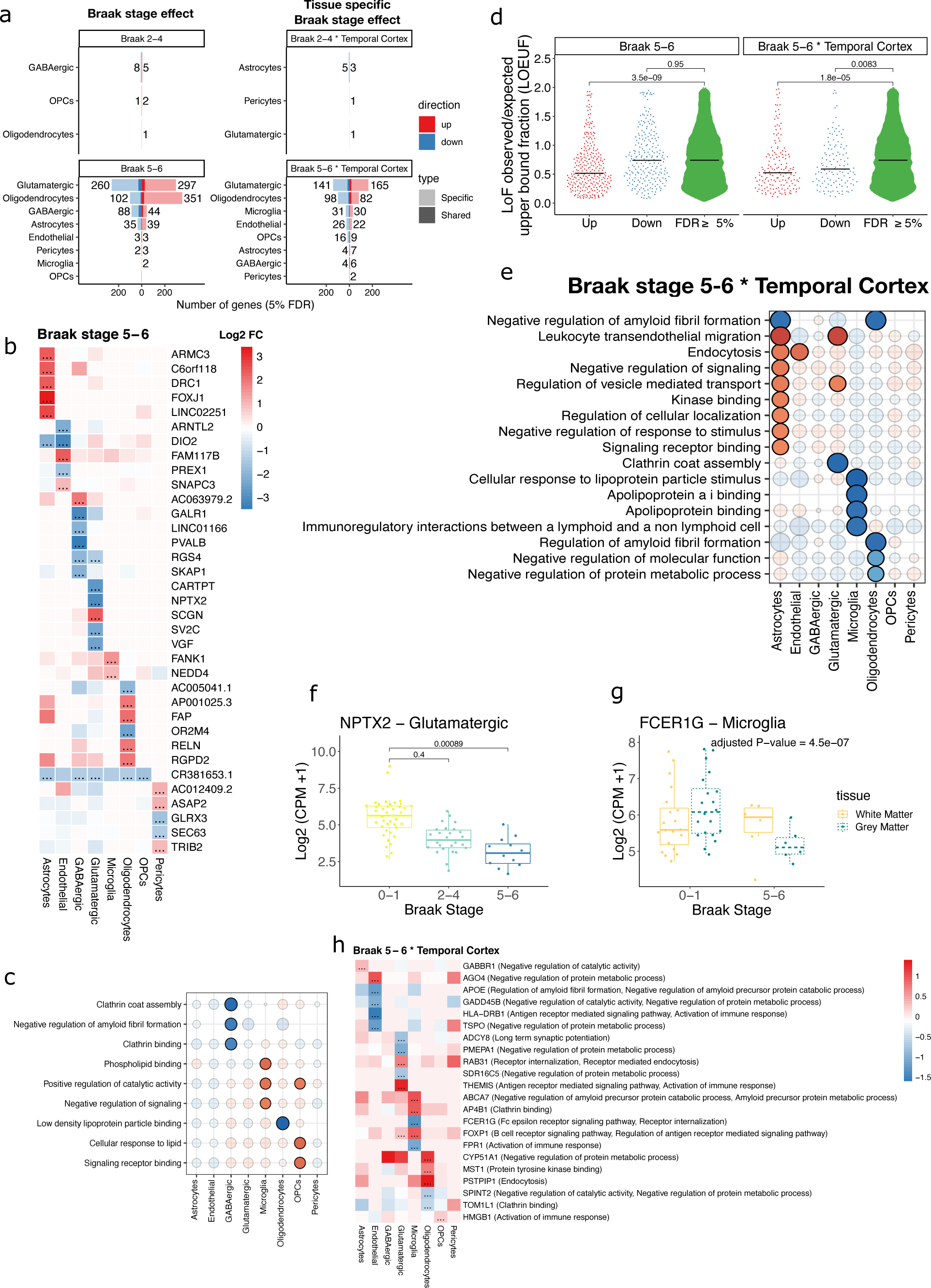
Differential expression analysis at cell type level. **(a)** Number of differentially expressed genes (5% FDR). **(b)** Differentially expressed genes with largest log fold change for each cell type at late Braak stage (5-6) vs early Braak stage (0-1). **(c)** Pathway enrichment of differentially expressed genes in AD genetically associated pathways. **(d)** LOEUF score (a measure of gene constraint) for differentially expressed genes in glutamatergic neurons. **(e)** Pathway enrichment (10% FDR) of differentially expressed genes in AD genetically associated pathways (20% FDR). **(f)** Expression of *NPTX2* in Glutamatergic neurons at different Braak stages. **(g)** Expression of *FCER1G* in Microglia at different Braak stages and in GM vs WM. **(h)** Differentially expressed genes affected in a different manner between GM AND WM that belong to AD genetically associated pathways (5% FDR).

However, we found a large number of differentially expressed genes (1230 at FDR-adjusted p-value <0.05) in late Braak stages (Braak 5-6 vs Braak 0-1), which is not unexpected given that cortical pathologies in the TC are only observed at later Braak stages in AD. For example, *NPTX2* (Fig.3f), a putative CSF prognostic AD biomarker^26^, was significantly lower in late Braak stages in glutamatergic neurons. Interestingly, the majority of differentially expressed genes were specific to individual cell types (∼90%, 1102/1230), suggesting that the cellular response to AD may not be broadly shared across cell types. We also found that 644 genes (FDR-adjusted p-value <0.05) were affected differently by late stage pathology in gray vs white matter (Fig.3a). For example, *FCER1G*, a gene associated with AD genetic risk^27^ was affected differentially in late Braak stages in gray (lower) vs white matter (higher) (Fig.3g), suggesting that transcriptional changes in AD can differ between different brain tissue types. Upregulated genes in late Braak stages (5-6) in glutamatergic neurons, as well as genes differentially affected between grey and white matter in glutamatergic neurons were significantly more constrained than non-differentially expressed genes (Fig.3d), suggesting that differentially expressed genes in glutamatergic neurons in AD also play an important developmental role. The differentially expressed genes (Braak 5-6 vs Braak 0-1) with the largest effect sizes included several interesting genes (Fig.3b). For example, *FOXJ1*, a key transcription factor involved in the production of motile cilia^28^, was significantly upregulated in Astrocytes. In addition, *PVALB* was significantly downregulated in inhibitory neurons, which is consistent with our proportion analysis (Fig.2a). We also found that *RELN*, which inhibits tau phosphorylation^29^, was strongly upregulated in oligodendrocytes.

Pathway enrichment analysis of these differential genes (Braak 5-6 vs Braak 0-1) highlighted several associated with Alzheimer’s disease (Methods). For example, genes involved in the negative regulation of amyloid fibril formation were lower at late Braak stages in inhibitory neurons, while ‘phospholipid binding’ and ‘cellular response to lipids’ were higher at late Braak stages in microglia and OPCs, respectively (Fig.3c). The genes with higher expression GM astrocytes in late stage pathology were enriched in endocytosis and kinase binding processes (Fig.3e), while the analogous genes in GM glutamatergic neurons were enriched in the regulation of vesicle transport. Genes with higher expression in WM microglia (but not GM microglia) at later Braak stages were enriched in cellular response to lipoprotein particle stimuli, while the analogous genes in WM oligodendrocytes showed enrichment for the regulation of amyloid beta formation. Finally, we looked for differentially expressed genes that were affected in a different manner between gray and white matter in late stage pathology and belonged to AD genetically associated pathways (Fig.3h). We found that *APOE*, a well-known AD risk gene belonging to pathways related to amyloid formation, is differentially expressed in endothelial cells between gray and white matter cells (with higher negative fold-change in GM as compared to WM). *ABCA7*, another AD risk gene^30^, was significantly higher in GM microglia than in WM microglia. Altogether, these results indicate that gene expression changes related to AD pathology are both cell type specific and tissue specific, suggesting that cross-tissue differential expression analysis at the cell type level will be necessary to fully capture the transcriptional changes associated with AD.

### Characterizing susceptible and resilient cell signatures with high-resolution spatial transcriptomics and pathological staining

To complement our analysis of dissociated nuclei from tissue, we next examined intact tissue using the CARTANA *probe-based in situ sequencing* platform with 155 pre-selected genes on 13 tissue samples from our donors^14^ (Supplemental Table1). In addition, we also performed staining on the same tissue sections to localize amyloid and tau pathological inclusions. The workflow included detection of β-amyloid and tau pathology, classification of 100×100 pixel (1056um^2) ‘tiles’ based on the quantitative plaque/tangle density, segmentation of white and gray matter, labeling of cell, and quantification of expression of 155 genes. This ultimately generated a resource with cells assigned to their major class, tissue coordinates of every pathological inclusion, and annotation of cells as present in white or gray matter (Methods, Fig.4a,b, Extended Data Fig.4a-c, Extended Data Fig.5a,b).

**Fig.4:**
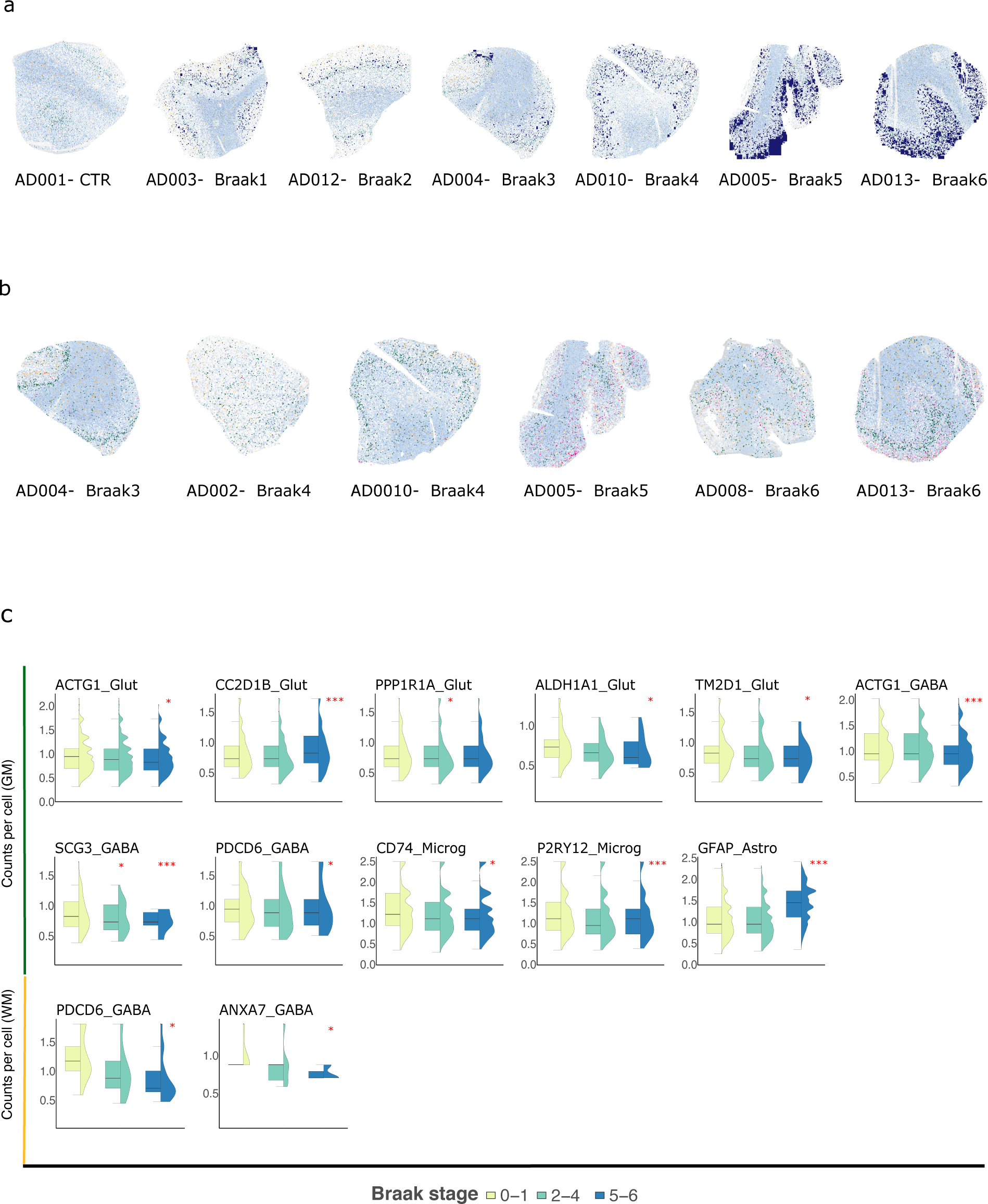
Characterization of plaque and tangle pathology in brain sections and validation of snRNA-seq derived associations via spatial analysis. **(a)** A visual representation of the spatial distribution of cells in the specified samples (from control to progressing AD pathology), with a background grid for all samples based on cell coordinates while different layers highlighting cells expressing specific markers (Oligodendrocytes =light blue to dark blue, *RORB*=green and *LAMP5*=orange). GM and WM are well distinguished with plaque deposition sites (blue squares) on tissue. **(b)** Same as ‘a’ except for tangles deposition instead of plaques in magenta squares. **(c)** Boxplots of the genes found to have statistically significant associations with Braak stages in GM and WM of TC on tissue validating snRNA trait association signatures. ‘*’ represents *p-value =<0.05* and ‘***’ represents *p-value =<0.01*.

Given the sparsity of detection in many of the genes, we were not able to definitively assign every cell to an snRNA-seq-derived subcluster, although the majority of cells did receive a ‘major class’ label (Supplemental Table1). Thus, instead of calculating cell subcluster counts directly in the tissue, we modeled the expression of subtype-specific genes in tissue by fitting a linear mixed model that investigated cell type-specific association of these marker genes and pathology (Supplemental Table1).

### Validation of snRNA-seq derived associations in neuronal and glial populations with advanced Braak stage

Our first analysis using the ISH data was to identify which snRNA-seq derived proportion differences were also present in intact tissue. Within glutamatergic neurons, we found that the expression of *ACTG1*, *ALDH1A1*, and *TM2D1* was statistically significantly lower in GM tissue from donors at late Braak stages, consistent with lower proportions of the *SPARCL1+* and RORB*+/ILRAPL2+* glutamatergic clusters in the dissociated nuclei (Fig.4c, Fig.2a, Extended Data Fig.4d, Supplemental Table1). Within GABAergic neurons, we found lower expression of *ACTG1* and *SCG3* which mark the *PVALB+/SPARCL+* subgroup in Braak stages 5/6 in GM, consistent with our snRNA-seq findings (Fig.4c, Supplemental Table1). In addition to this, we also found relatively lower expression of *PDCD6* in Braak stages 5/6, consistent with our GABAergic *PVALB+/TMEM132C+* subgroup in GM. Finally, our finding on the relatively resilient *THEMIS+/POSTN+* glutamatergic population was also confirmed in our ISH data, where we found higher expression of *PPP1R1A* and *CC2D1B* within glutamatergic neurons at Braak stages 2/4 and 5/6, respectively (Fig.4c, Extended Data Fig.4d, Supplemental Table1).

On the glial side in GM, the ISH data recapitulated lower proportions of the *RPL19+* microglial subcluster, as evidenced by lower expression of *CD74* and *P2RY12* in microglia in intact tissue (Fig.4c, Supplemental Table1). By contrast, higher expression of *GFAP* at advanced Braak stages was consistent with our snRNA-seq findings regarding higher proportions of the *TNC+ (GFAP+)* astrocyte cluster at Braak stage 5/6 (Fig.4c, Extended Data Fig.4e, Supplemental Table1).

With respect to WM cell signatures, we identified lower expression of *PDCD6* and *ANXA7* in *PVALB+* GABAergic neurons in Braak stage 5/6, corresponding to our earlier finding of lower proportions of PVALB+/TMEM132C+ and *PVALB+/SPARCL1+* GABAergic clusters (Fig.4d, Supplemental Table1). Thus, even though neuronal cells are rare in white matter, our findings for *PVALB+* subgroups are consistent across the dissociated nuclei and intact tissue.

In addition to validating gene expression with each broad cell class in our *in-situ* data, we also examined global signatures aggregated across all cell classes in both GM and WM (Extended Data Fig.6a) at different Braak stages. At the global level, we did not find any statistically significant genes after multiple testing corrections. However, we found 29 genes with nominal significance, which is a two-fold enrichment with respect to the null. For example, we recapitulated our prior astrocyte-specific analysis and found a general increase in *GFAP* expression at late Braak stages 5/6. At the case-control level, we identified 31 genes with a nominal p-value below 0.05, which is a ∼ 4-fold enrichment compared to what would be expected under the null (i.e., 155 tests*0.05=8 genes) (Extended Data Fig.6.b). Notably, we found a decrease in *SST* expression (significantly downregulated in GABAergic neurons in our single nucleus data), as well as a decrease in *NEFL* expression (significantly downregulated in excitatory neurons in our single nuclei data), confirming their downregulation in AD. Thus, at both the global and broad cell class level, our *in-situ* analysis confirmed some of our strongest signals from our dissociated single-nucleus RNA-seq proportion and gene expression analyses with nominal statistical significance.

### Spatial analysis identifies genes with altered expression near pathological inclusions

We next took advantage of the spatial information afforded by our multiplexed ISH approach to identify cell type-specific gene expression changes in the microenvironment of plaques and tangles in GM tissue. We used two approaches - a distance-based model and a tile-based model - to examine differences in tissue composition near and far from plaques/tangles.

For the distance-based model, we categorized every cell into one of three groups: close (< 70 μm) to plaques, intermediate distance from plaques (70 - 154μm), and far away (>154 μm) from plaques and then identified statistically significant differences in cell type-specific gene expression in cells belonging to each of these three groups. In glutamatergic cells, there was higher expression of *ALDH5A1* and *THEMIS* expression in the plaque-distant group, whereas *NEFL* and *NEFM* showed lower expression in this group (Fig.5b). In GABAergic cells, *RELN* showed higher expression in the plaque-intermediate and distant group, consistent with previous findings that *RELN+* interneurons are likely affected in AD^31,32^. In microglia, expression of *CD68* was higher in plaque-intermediate distance cells. In astrocytes, *GFAP* expression was higher in plaque-distant cells. Finally, in endothelial and pericytes, *ID3* showed higher expression in plaque-distant cells (Fig.5b).

**Fig.5:**
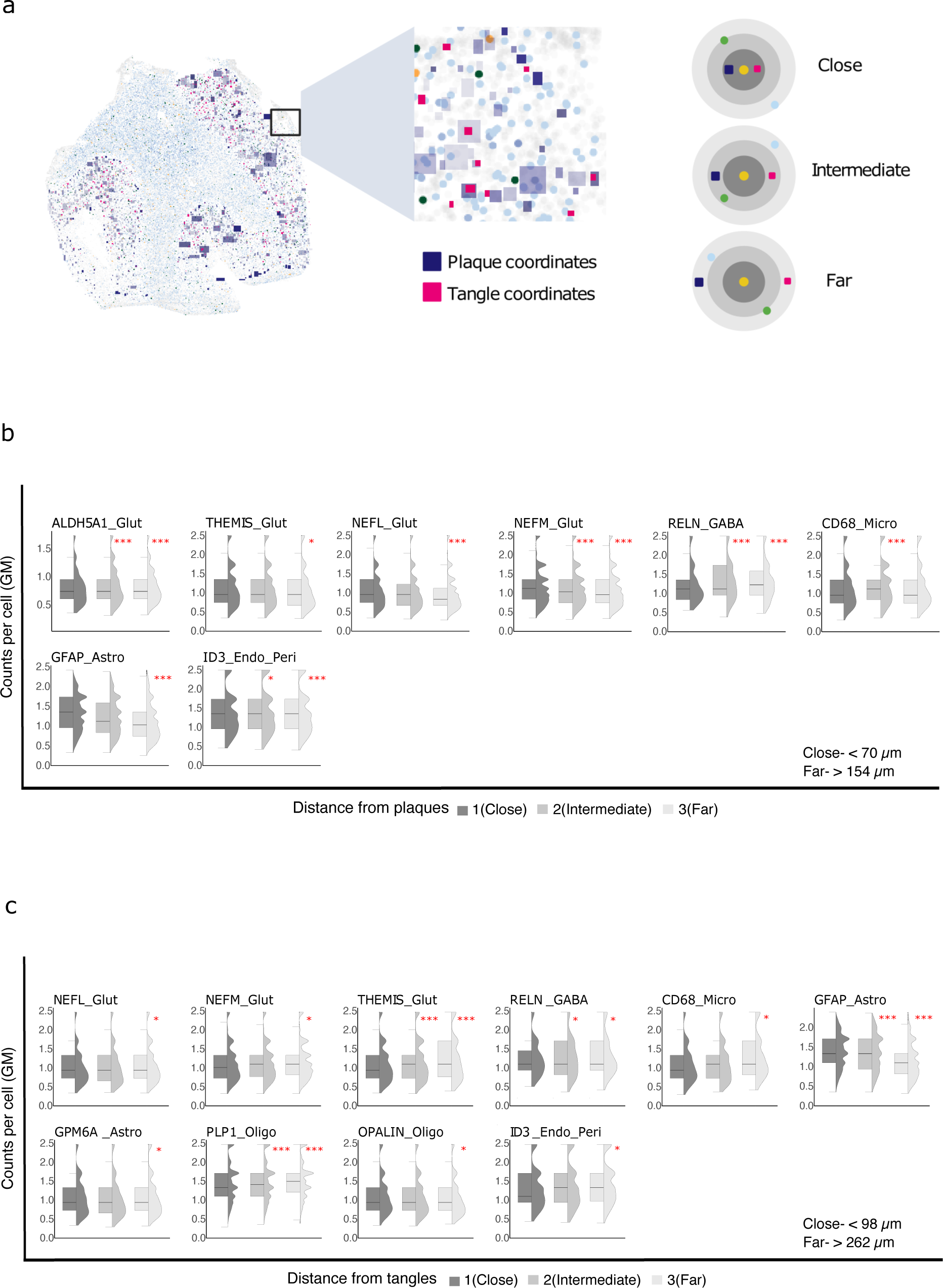
Gene expression in tissue niches of plaques and tangles. **(a)** A cartoon representing the distance for a cell to get assigned as close, intermediate, or far from plaque/tangle depositions. **(b)** Boxplots of the genes found to have statistically significant associations with distance from plaques in GM of TC on tissue. **(c)** Same as ‘b’, showing statistically significant associations with distance from tangles. ‘*’ represents *p-value =<0.05* and ‘***’ represents *p-value =<0.01*. Note that for *GFAP* the visualization does not reflect the increase in *GFAP* expression in pathology distant cells (as found in the model); this discrepancy may result from the lack of accounting for donors in the visualization.

For the tile-based analysis, we divided the tissue into 100 square micron regions, and identified those that contained amyloid plaques versus those that did not. This alternative model takes into account not just the closest plaque, but rather captures differences due to the overall plaque density in a given microenvironment. Although this analysis did not yield any significant genes after multiple testing correction, 15 genes were nominally significant (p<0.05), corresponding to a 2-fold enrichment with respect to the null hypothesis (Extended Data Fig.6c). Similar to our distance-based analysis, this approach also detected an increase in *NEFM* expression in glutamatergic neurons in plaque-containing tiles. In microglia, we observed an increase of *BIN1*, a well known AD risk gene, in plaque-containing tiles. Finally, we found a strong reduction of *SERPINE1*, a vascular and microglial-specific gene involved in the inhibition of plasmin which degrades A-beta plaques, in close proximity of plaques only in individuals with low pathology, suggesting that activation and abeta degradation may be greater in these individuals (Extended Data Fig.6c).

We also repeated our cell type-specific gene expression analysis based on proximity to tangles in GM tissue and observed more complex relationships. Given the higher density of tangles compared to plaques (in individuals with substantial tangles), we restricted our analysis only to the distance-based model. With this model, we found higher expression of *NEFL, NEFM* and *THEMIS* in the tangle-distant group (>262μm) compared to tangle-proximal group (<98μm) (Fig.5c). In GABAergic neurons, *RELN* had a higher expression in the tangle-intermediate and distant group, consistent with the plaque-proximal finding presented above (Fig.5c). Microglial *CD68* gene expression differences associated with distance from tangles also reflected higher expression in the tangle-distant group (Fig.5c). *GPM6A* expression in astrocytes and *PLP1* along with *OPALIN* were higher in oligodendrocytes in tangle-distant cells. *GFAP* expression in astrocytes also showed a similar spatial pattern with respect to tangles as with plaques, increasing in tangle-distant group. Finally, *ID3* expression was higher in tangle-distant endothelial and pericytes group (Fig.5c). Overall, this suggests a set of coordinated changes across multiple glial cell types in the local environment surrounding tangle-bearing neurons, with many distributional changes shared between extracellular plaques and tangles.

## Discussion

Our study is among the first to investigate transcriptomic differences in the temporal cortex in controls and AD donors by employing two techniques; snRNA-Seq and *ISH* with immunohistochemical characterization of pathology using the CARTANA platform. We report a total of 430,271 single-nuc transcriptomes across 80 individuals (40 in GM and WM each) with a range of AD-associated pathology as well controls. Our overall findings can be grouped into four sets of discoveries, as summarized below.

First, we found that multiple subpopulations of neurons had lower proportions in GM in donors at late Braak stages, with the exception of glutamatergic subpopulations marked by *THEMIS* and *POSTN. THEMIS*, previously described in T cells, has been identified as a marker of deep layer neurons in humans, and periostin (*POSTN)* is an extracellular glycoprotein originally identified as a molecule differentially expressed in osteoblasts and fibroblasts^33^. Recent studies have shown that *POSTN* exhibits neurite outgrowth activity in cerebellar granule neuron (CGN) or dorsal root ganglion (DRG) neurons^34^ and neuroprotective activity^35^ in adult cortical neurons^36^. This suggests a potential neuroprotective role of *POSTN* in neurons that are less affected by AD pathology.

With respect to GABAergic neurons, the higher proportion of *TRHDE*-expressing cells at later Braak stages suggests that thyrotropin-associated pathways may show differential regulation in disease. The other GABAergic neuron marker gene with altered expression in individuals with pathology is *THSD7B,* which has been reported to be associated with age-related cognitive decline based on repeated measures of 17 cognitive tests suggesting that this GABAergic subpopulation could also serve as a candidate for future perturbation studies^38^.

Second, we found multiple glial subgroups with altered proportions in AD, including *RPL19+* microglia and *TNC+* astrocytes. The former show lower proportions in tissue from individuals at late Braak stages, whereas the latter shows the opposite effect. However, our cross-sectional study does not provide information as to whether these signatures are protective, reactionary, or pathological. Indeed, prior work on the extracellular matrix protein *TNC* (Tenascin) has shown it to be at very low levels in the adult nervous system and found to be upregulated in lesioned adult mouse brain^39^. In human brain tissue, *TNC* has also been shown to have high expression in EC and SFG within *GFAP* high astrocytes^10^.

Third, we found surprising differences in neuronal composition in WM, despite the overall rarity of neurons in this tissue. The *RORB+/ILRAPL2+* glutamatergic neuron subgroup shows higher proportions in WM in intermediate Braak stages, in contrast to the lower proportions in GM at late Braak stages, suggesting either potential mislocalization or differences in pathological processes in GM and WM. By contrast, we found that *LAMP5+/CHST9+,* PVALB+/TMEM132C+ and *PVALB+/SPARCL1+* GABAergic clusters are preferentially lower in WM at late Braak stages. Thus, even though neuronal cells are rare in WM, we identified neuronal distribution changes in WM in both directions, which were also validated in our multiplexed ISH data. Interestingly, we also observed some sex specific enriched/depleted sub-populations in TC and multiple regions as well (Fig.2e, Extended Data Fig.3c-f). Despite our inability to confirm the correlation of these signatures with sex, it’s prudent to catalog them for potential future interventions, particularly in light of their possible AD specificity.

Fourth, our multiplexed ISH data, combined with immunostaining for pathology on the same section, allowed us to characterize cell type-specific gene expression differences between regions close to and far away from amyloid plaques and neurofibrillary tangles. Given differences in localization of plaques (extracellularly) versus neurofibrillary tangles (primarily within neurons), it is not surprising that the pathology-proximal signatures of neurons were different in the two cases. This could reflect selective vulnerability of neuronal classes described in the Results section to each of the two pathologies. By contrast, microglial signatures in both plaque-proximal and tangle-proximal regions showed consistently lower expression of genes such as *CD68* and astrocytes in both pathological microenvironments also showed shared signatures, including higher expression of *GFAP* in low pathology environments. Thus, by identifying cell type-specific genes that are likely altered in proximity to AD proteinopathies, we highlight that there are coordinated changes across almost all major cell types in space, which may not be observable in studies looking at dissociated nuclei. Although we cannot establish causality from proximity-based studies such as this one, we can nonetheless refine the set of candidate signatures that are most likely to be interacting directly with pathological inclusions in AD.

Given the novelty of the combined methods described here, it is important to contextualize our findings in light of some of the limitations of the experimental approaches. Notably, our use of multiplexed ISH through the CARTANA platform does suffer from sparsity, leading to non-comprehensive assignment of individual cells to broad cell classes or subtypes. Although our cell class-specific differential expression analysis did allow us to replicate a subset of our snRNA-seq findings, several of the subgroups identified through snRNA-seq were not clearly delineated in ISH, preventing the possibility of validating their proportion differences. In addition, our cohort size (40 individuals), while substantial, is smaller than those from recent studies on the prefrontal cortex^31,40^, suggesting again that there may be some false negatives masked by the heterogeneous presentation of AD and the need for larger cohort studies.

Overall, the study presented here provides a multimodal examination of GM and WM in temporal cortex, examining both compositional changes as well as spatial differences in tissue from individuals at various Braak stages. Together with other studies looking at cell type differences in post-mortem tissue from other brain regions (primarily GM), this type of examination is necessary not only to characterize cell type-specific alterations in AD, but also to refine a set of candidate signatures to probe in targeted downstream experiments. As with prior studies, we find that multiple classes of cells in the brain show differences in individuals at early versus late stages of pathological accumulation, highlighting key compositional and spatial alterations that may ultimately lead to novel cellular signature-based therapeutics.

## Methods

### Nuclei isolation and library preparation for Single-nucleus RNA-Seq

Nuclei were isolated from 10-μm fresh-frozen sections using the Nuclei PURE Prep Nuclei Isolation Kit (Sigma-Aldrich), with specific modifications. Regions of interest were macro-dissected using a scalpel blade and lysed in Nuclei Pure Lysis Solution containing 0.1% Triton X, 1 mM DTT, and 0.4 U µl−1 SUPERase-In RNase Inhibitor (Thermo Fisher Scientific), freshly added before use. The samples were homogenized sequentially with 23-gauge and 29-gauge syringes.

Cold 1.8 M Sucrose Cushion Solution, prepared immediately before use with 1 mM DTT and 0.4 U µl−1 SUPERase-In RNase Inhibitor, was added to the suspensions, which were then filtered through a 30-μm strainer. The lysates were carefully layered on top of 1.8 M Sucrose Cushion Solution in new Eppendorf tubes and centrifuged for 45 minutes at 16,000g and 4 °C. Resulting pellets were resuspended in Nuclei Storage Buffer containing 0.4 U µl−1 SUPERase-In RNase Inhibitor and transferred to new Eppendorf tubes for centrifugation at 500g and 4 °C for 5 minutes. This step was repeated once.

Finally, purified nuclei were re-suspended in Nuclei Storage Buffer with 0.4 U µl−1 SUPERase-In RNase Inhibitor, stained using trypan blue, and counted with Countess II (Life Technologies). Approximately 12,000 cells per sample were loaded onto the 10x Single Cell Next GEM G Chip. cDNA libraries were prepared using the Chromium Single Cell 3′ Library and Gel Bead v3 kit, following the manufacturer’s instructions. Sequencing was performed on the Illumina NovaSeq 6000 system with the NovaSeq 6000 S2 Reagent Kit version 1.5 (100 cycles), targeting a minimum sequencing depth of 30,000 reads per nucleus.

### Data processing and quality control

All samples from our temporal cortex dataset, as well as from the public datasets, underwent uniform processing using Cell Ranger (version 3.1.0), employing the GRCh38 reference human genome and the Ensembl Homo_sapiens GRCh38.91 reference annotation, modified to include intronic reads. Nuclei were characterized as barcodes containing a minimum of 500 unique molecular identifiers (UMIs) (excluding mitochondrial RNA) and less than 5% mitochondrial RNA. For samples with over 10,000 nuclei, only the top 10,000 nuclei with the highest UMI counts were retained. Doublets were marked using scDblFinder^41^ (version 1.4.0), but not excluded as none of the clusters were enriched in doublets. We filtered out two samples (from one individual) due to low quality (less than 500 cells with a lower amount of UMI, detected genes and higher mitochondrial reads percentage than the other samples) (Extended Data Fig.1e). An overview of the TC region study including the number of samples from varying pathology and controls are depicted in Extended Data Fig.1a-b. The 80 samples from TC region were integrated and clustered using Conos^42^ (Extended Data Fig.1c) and Harmony^43^ (Extended Data Fig.1d) and based on our observation with the integrated data from multiple brain regions (Extended Data Fig.2 c-f) and its cell type specific sub-clustering (Extended Data Fig.3 a-b) we considered Harmony as our final integration method for TC region as well as multiple brain regions. The primary QC visualization for TC region and multiple published studies can be seen in Extended Data Fig.1a-b. We used ‘*findMarkers*’ from the scran R package to identify markers for the different cell types and cell states in the TC region^44^. More specifically, we used the *‘multiMarkerStats’* function which combines the results of three different statistical tests (wilcox, t-test and binomial test).

### Integration of multiple datasets from multiple brain regions

We integrated our dataset from TC region (GM and WM) to other published studies from different brain regions that include entorhinal cortex^10,11^ (Leng et al^8^, Grubman et al^9^), prefrontal cortex^12,13,5^ (Cain et al^12^, Zhou et al^13^, Mathys et al^5^), superior frontal gyrus^10^ (Leng et al^10^) and deep white matter from prefrontal cortex^12^ (Cain et al^12^). The matrix of filtered UMI counts (>500) for each study was converted to a single cell experiment (SCE) object using the function *read10xCounts* from SingleCellExperiment^45^ package on R (version 4.0.1) platform. The data class conversion from *sce* to Seurat object and merging of all datasets were executed using conventional functions *CreateSeuratObject* and *merge* respectively from Seurat package (Version 3.2.0.)^46^. The merging of all datasets did not colocalize the same brain regions (Extended Data Fig.2c) that led to the important step of integration. We integrated all datasets using the reciprocal PCA approach which was implemented within the Seurat package that identifies anchors utilizing reciprocal PCA (RPCA) instead of canonical correlation analysis (CCA) (Extended Data Fig.2e). We also used Harmony^43^ (version 0.1.0) within the Seurat workflow using the *RunHarmony* function (Extended Data Fig.2f) to integrate all datasets. Other than these two approaches we also used Conos^42^ (version 1.4.6) to integrate multiple brain regions’ datasets from aforementioned published studies (Extended Data Fig.2d). Before performing the integration, we followed the conservative steps of normalizing, scaling the counts and running the PCA on a total of 959,237 nuclei using inbuilt functions from the Seurat package and selected 30 principal components for the Harmony and RPCA integration, while we used 50 principal components for the Conos integration. The Conos (version 1.4.6) (resolution = 0.8) and Harmony (version 0.1.0) (resolution = 0.8, ‘study’ as the integration variable) integration were subsequently labeled independently based on canonical marker expression. A subset of small clusters from the Conos integration, expressing multiple canonical markers, were designated as ‘mixed’. Nuclei not labeled as ‘mixed’ received Conos cell type annotations, while the remaining nuclei were assigned Harmony cell type labels, though we observed a good agreement of Conos and Harmony for the integration of the major cell (Extended Data Fig.2i). Since the Harmony algorithm iteratively corrects PCA embeddings, the downstream analyses used the Harmony embeddings instead of PCA. We clustered the integrated datasets using *Seurat::FindClusters* function at a resolution of 0.8, obtained from both approaches. This function clusters the cells using a shared nearest neighbor (SNN) modularity optimization approach. These pre-clusters were annotated using a canonical set of marker genes and were split into 8 broad cell type clusters (Extended Data Fig.2g, Supplemental Table1) with a reasonable distribution of various cell types (Extended Data Fig.2h) that were visualized employing a 2D UMAP projection executing *DimPlot* function from Seurat package and a bar plot using *dittoSeq::dittoBarPlot* function^47^ respectively.

### Sub clustering of major cell types within the integrated dataset

After identification and clustering of broad cell types at the top level, we sub-clustered all broad cell types independently using RPCA and Harmony to investigate the further subtypes within each class. To subcluster, we started from sub-setting the specific cell type cluster from the integrated dataset and reintegrated them as aforementioned. We compared cell type specific reintegration and subclustering and noticed Harmony to integrate the data from the same brain region more relevant (Extended Data Fig.3a,b). Specifically, OPCs and endothelial cells integrated by RPCA did not integrate the PFC region datasets reasonably (Extended Data Fig.3a). The downstream analyses were performed on subclusters generated from Harmony (Supplemental Table1) as the consistency of the subclusters from the same brain region in the integrated dataset were more accurate with this approach.

### Machine learning based refinement of integrated clusters

We used Random Forest as a robustness method for our subclusters obtained for all major cell types. To train the model, we included the subclusters within each cell type with at least 50 cells where 75% of the cells could be used as a training set and remaining 25% for testing at minimum with topmost variable genes. We identified genes that drove biological heterogeneity in the data set by modeling the per-gene variance using the function *modelGeneVar* from Bioconductor package scran^44^ (version-1.18.7). After predicting the classes on the testing set of clusters, we evaluated the accuracy of a classification with the help of a confusion matrix (Extended Data Fig.7). We merged the subclusters based on their higher similarities produced by the random forest classification.

### Cross-comparison between Integrated and TC region specific clusters

We compared the clusters from the TC region to the integrated ones and ran another round of refinement of clusters. This led us to merge some additional clusters in the integrated dataset for each broad cell type (Extended Data Fig.8). The integrated clusters were annotated based on top cluster markers present within each identity cluster. We obtained these markers using the function *FindAllMarkers* from the Seurat package. We only tested genes that were detected in minimum 25% cells in a subcluster and showed at least 0.25-fold difference (log-scale). The glutamatergic subclusters were annotated using layer marker genes co-expressed with other top cluster identity markers. The layer association of top gene markers for glutamatergic cell types were established using transcriptomics explorer offered by Allen brain map atlas encompassing human multiple cortical areas (*<u>Transcriptomics Explorer :: Allen Brain Atlas: Cell Types</u>.)* Our multiple layered refinement approach led us to have 888,784 nuclei to analyze further. (Table1, Extended Data Fig.2j).

### Trait association in TC region and comparison to different brain regions

We removed two bad quality samples, and all dementia samples from our investigation of cell proportion changes with the AD pathology in the TC region. To explore the cluster proportion changes in early or late AD pathology in WM and GM of TC region we implemented ANCOM-BC^48^ (Analysis of Compositions of Microbiomes with Bias Correction, version_1.0.5) which is a log-linear model that determines differentially abundant taxa according to the variable of interest. We inspected WM and GM from TC in each cell type independently to investigate any associations of the changes in cluster proportion with AD pathology. The clusters were implemented as taxa and the AD pathology marked by different Braak stages was used as a covariate of interest. We regressed out sex, age of death and postmortem interval. The numeric covariates like age of death and postmortem interval were divided by two times their standard deviations to scale the regression^49^. The Braak stages were classified in three discrete categories. The Braak stages less than 2, ranging from 2-4 and more than 5 were designated as ‘Braak_disc 0-1’, ‘Braak_disc 2-4’, and ‘Braak_disc 5-6’ respectively.

We implemented the same strategy for the integrated cell type classes with an additional covariate of different brain regions in the model. Before fitting the log linear model, we removed any cells from the integrated dataset that had no information of PMI, age of death, sex, and Braak stage. We also excluded the *Grubman et al* study from this part of analysis due to discrepancy in demographic details. This additional filtering resulted in a total of 835,413 nuclei for the trait association analysis that were derived from a total of 216 samples. The trait association analysis was performed separately for each broad cell type.

### Differential expression analysis

We performed our differential expression analyses using a ‘pseudo-bulk’ approach, that is we summed the total number of UMI for each sample and each cell type, resulting in a single count value for each gene in each sample and each cell type. We only retained samples which had at least 10 nuclei from a specific cell type, and sub-populations that were detected in at least 20 samples. We also only tested genes with a median count per million (CPM) of at least 1 and expressed in at least 20 samples. Furthermore, we removed genes with low counts using the *‘filterByExpr’* function from the *edgeR*^50^ R package. We then used the *glmmTMB* R package^51^ to fit a negative binomial mixed model to the data for each cell type (family = nbinom2). We used the following models: counts (raw) ∼ Braak_stage (or diagnosis) * tissue (gray/white) + age + sex + pmi + (1|individual) (random effect as each individual has a sample from gray/white matter), effectively testing for an effect of Braak_stage (or diagnosis) in our cohort, as well as allowing brain region specific effects. As an offset, we used the log of the TMM normalized library size^52^ minus the log of 1 million. The following variables were categorized prior to model fitting as this improved the convergence of our statistical model: Braak_stage (0-1, 2-4, 5-6), age (<60, 60-80, >80), pmi (<6,6-8,>8).

### Enrichment of differentially expressed genes

First, we identified genetically enriched AD pathways, by using MAGMA^53^. Briefly, we obtained gene-level genetic associations using GWAS summary statistics from a recent AD GWAS^54^ (using all SNPs within a window starting from 35kb upstream of each gene to 10kb downstream). We then used MAGMA to test whether any pathways from the GO_BP, GO_CC, GO_MF, Hallmark, KEGG and REACTOME databases were enriched in AD genetic risk. Multiple testing correction was performed using the Benjamini-Hochberg procedure. Finally, we then tested whether the differentially expressed genes were enriched in any AD genetically associated pathways using the fgsea^55^ R package.

### In-situ sequencing

#### Immunostaining of human samples

Based on previously published studies we chose 155 genes (Supplementary Table1) to analyze the spatial arrangement of cell types on the tissue sections of 13 individuals comprising different stages of AD pathology as well as controls. We used CARTANA to generate and sequence clonally amplified barcode sequences deploying gene specific probes ligation onto tissue sections.

Neuropathologically characterized human brain blocks from controls and Alzheimer’s patients obtained from Netherland Brain Bank were cryosectioned into 10µm sections, onto SuperFrost Plus glass slides (ThermoFisher) and stored at −80°C. In situ sequencing was performed using the reagents supplied by CARTANA^14^ (PN1110-02, HS Library Preparation Kit; PN3110-02 - In situ sequencing kit) and following the recommended protocol for human fresh frozen samples, with autofluorescence quenching. Briefly, the sections underwent fixation, permeabilization, probes hybridization and ligation, rolling circle amplification, and seven sequential cycles of fluorescence labeling and image acquisition at 20x magnification on an ECLIPSE Ti2 inverted microscope (Nikon) (Extended Data Fig.4a). The probes used belonged to a panel of 155 genes, designed for this study by CARTANA, based on their state-of-the-art guidelines.

After completing in situ sequencing, the same sections underwent immunostaining of amyloid plaques and phosphorylated Tau. Briefly, after removing the coverslips, in situ sequencing probes were detached through sequential washings with formamide. Afterwards, blocking solution (SP-5030-250, Vector Labs) was added for 20 minutes before performing an incubation with mouse anti-Phospho-Tau (Ser202, Thr205) (MN1020, ThermoFisher) for two hours, followed by one hour incubation with donkey anti-mouse IgG [H+L]/ AF 550 (SA5-10167, Invitrogen) and one hour incubation with mouse anti-Beta-Amyloid MOAB-2/AF 750 (NBP2-13075AF750, Novus Biologicals). Finally, sections were incubated for 30 seconds in 1x TrueBlack (23007, Biotium) to reduce autofluorescence and for 5 minutes in DAPI (62248, ThermoFisher) for nuclei detection. Sections were then mounted using SlowFade™ Gold Antifade Mountant (S36940, ThermoFisher) and imaging was carried out using a 20x magnification on an ECLIPSE Ti2 inverted microscope (Extended Data Fig.5a).

#### Gene expression

For each sample we obtained from in situ sequencing a reference DAPI image and 6 sequencing images (one for each barcode position) that were aligned and combined using ISSanalysis software (Quantified Biology) in order to generate a map of the spatial coordinates of each cell (Supplementary Table1) and each target RNA molecule on the section. Afterwards, for each tissue section, it was determined the total count of spots for each target gene and this value was normalized on the tissue area (Supplementary Table1). To compare the gene expression of different samples while removing any bias due to technical factors, the expression of each gene was normalized also on the total number of identified molecules per sample (Supplementary Table1).

#### Gene enrichment in plaques proximity

First Halo v3.1 (Indica Lab, https://indicalab.com/halo/) was used to build a classifier able to automatically recognize beta amyloid plaques from the immunostaining images (Extended Data Fig.4b). Plaques coordinates were then exported and uploaded on ISSanaysis software, where they were used to identify genes, whose expression was enriched in proximity of plaques. In order to do so, a grid was applied to each tissue section and for each tile was determined the presence/absence of a plaque and the presence of any of the *in-situ* sequencing target genes (Extended Data Fig.4c). Similarly, to what was done previously, for each sample it was determined the normalized count of reads for each gene in the areas with and without plaques (Supplementary Table1).

#### In situ cell-type classification

We used the following strategy to label cells in our CARTANA ISS experiment. First, we aggregated all counts for each gene and each broad cell type (across samples) from our temporal cortex single nuclei experiment. Second, we normalized the broad cell type pseudo-bulk expression data so that each broad cell type had a total of 10k reads (CP10k). We then computed a specificity metric for each gene by dividing the CP10k from each broad cell type by the total CP10k of all broad cell types. We then identified broad cell-type marker genes as genes with a CP10k of at least 0.01 and a specificity metric greater than 0.5 (indicating that more than 50% of the expression of the gene across cell types is quantified in the given cell type). We then used the subset of marker genes that were also quantified in our CARTANA ISH experiment to label cell types. For each CARTANA cell, we summed the total number of CARTANA counts across all markers from the different broad cell types (i.e., Astrocytes, Oligodendrocytes, etc.). Then, for each cell, we computed a z-score of the counts in marker genes from the different cell types, which were then transformed into a one-sided p-value using the *pnorm* R function. Finally, cell type labels (i.e., Astrocytes, Oligodendrocytes, etc.) were given to each individual cell from the tissue section if the p-value was below 0.05 (indicating that the sum of count for a marker of a specific cell type was significantly higher than counts of marker genes for the other cell types).

#### Pathology-induced gene expression variability in the spatial environment

We modeled the expression of 155 genes within each broad cell type for WM and GM of TC region separately to test whether there’s a significant association between the expression of 155 CARTANA genes and AD phenotypic trait (Braak stage). We fit a linear mixed-effects model to investigate this relationship of gene expression with different Braak stages while taking individual variation as random effect for each major cell type. Similar to our snRNA trait association analysis we categorized the Braak stages into three groups; Braak stage 0-1, 2-4 and 5-6 and modeled the association for every cell type in WM and GM of TC (Supplementary Table1).

#### Microenvironment -induced gene expression variability in the spatial environment

We modeled the expression of 155 genes based on the microenvironment of plaques and tangles depositions. Based on the cell coordinates of the plaques and tangles (Supplementary Table1) on the tissue we were able to represent the potential location of plaques and tangles for all the samples except for sample AD006 (Fig.4a,b). We computed the Euclidean distance of each cell from each plaque location for every sample and considered the minimum distance to the closest plaque for each cell to consider the microenvironment. The same strategy was applied to compute the tangles’ microenvironment. To associate the cell location from the plaques on the tissue we distributed the distances into three bins marking each cell either being close (bin1), intermediate (bin2) or far away (bin3) from the plaques or tangles. We then fit the linear mixed-effects model to investigate this relationship while including distance bin as the fixed effect while regressing out the variation coming from individuals (Supplementary Table1).

---

**Extended Data Fig.1:**
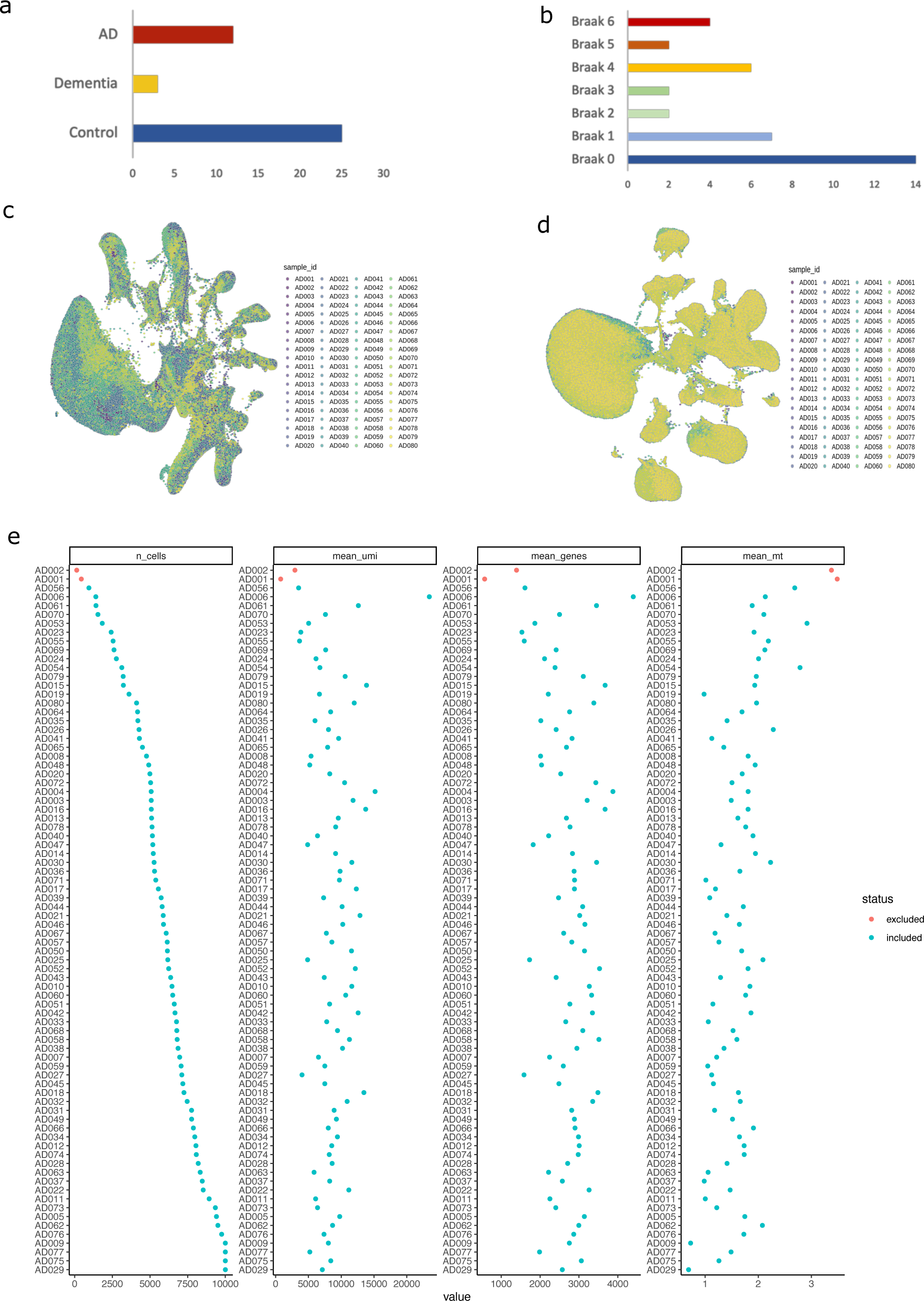
A cellular architecture of the human TC in 40 individuals: Quality controls. **(a)** Clinical information of individuals. **(b)** Barplot showing the number of different pathologies (varying Braak stages) for individuals. **(c)** UMAP embedding of 463,988 single-nucleus RNA profiles (pre-QC) from the TC brain region of 40 individuals including gray and white matter samples, generated by Conos integration method (80 samples). **(d)** same as ‘c’ instead of UMAP embeddings are generated by Harmony integration method. **(e)** QC plot for TC samples.

**Extended Data Fig.2:**
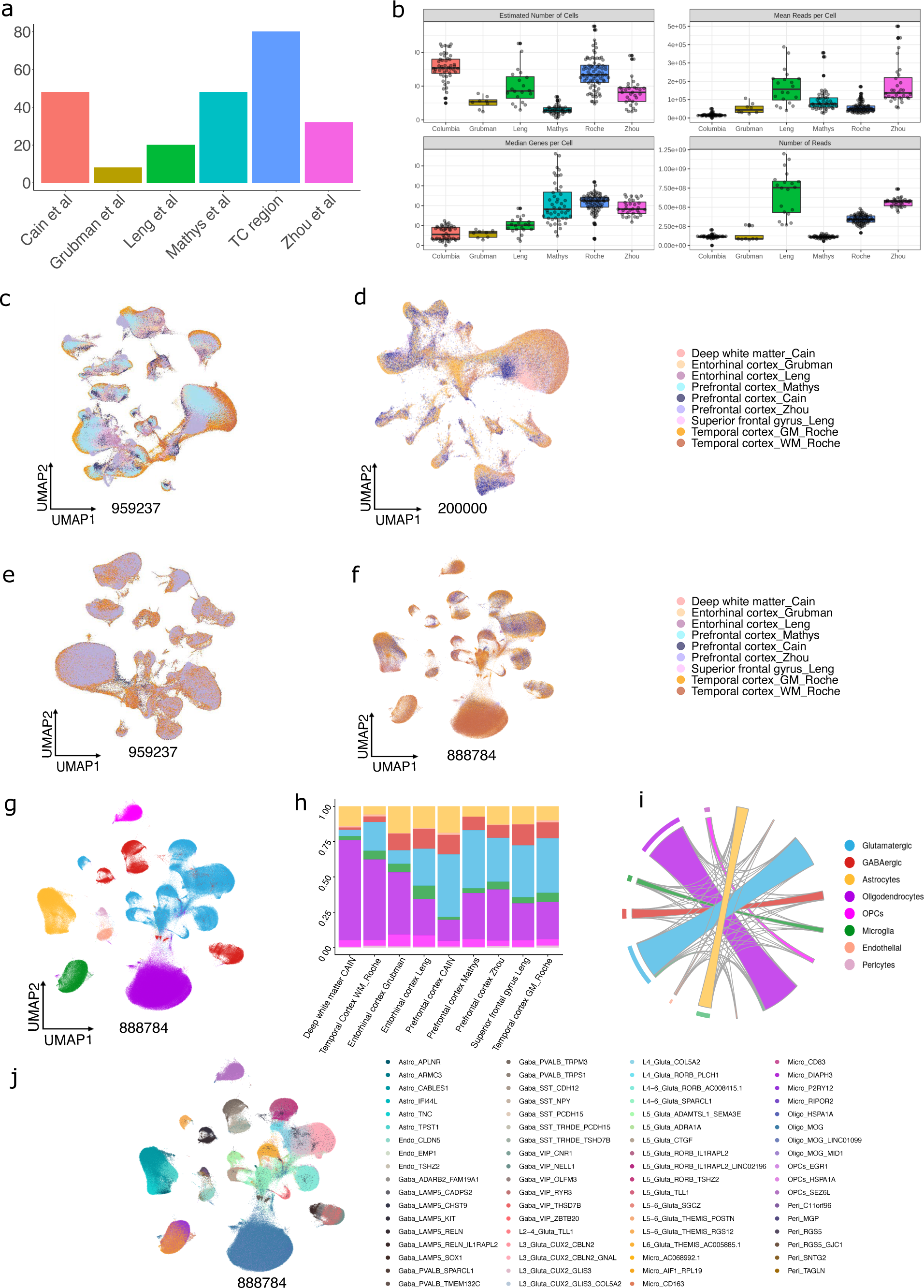
A cellular map of the multiple human brain regions. **(a)** Bar plot showing number of samples (y-axis) per study included for multiple brain region analyses. **(b)** QC plots for all studies/brain regions. **(c)** UMAP embeddings of 959,237 (pre-QC) nuclei after merging multiple brain regions. **(d)** UMAP embeddings of ∼200k nuclei after integrating multiple brain regions using Conos. **(e)** UMAP embeddings of 959,237 nuclei after integrating multiple brain regions using RPCA. **(f)** UMAP embeddings of 888,784 (post-QC) nuclei obtained after integration and additional filtering cells in multiple brain regions using Harmony. **(g)** UMAP (Harmony generated) embeddings of 888,784 nuclei colored by 8 major cell types. **(h)** Percent compositions of major cell types within each study. **(i)** Circular plot showing agreement between Conos and Harmony generated cell type annotations at the top level (peripheral annuli representing Conos labels). **(j)** subclusters for each cell type across multiple brain regions (888,784 nuclei).

**Extended Data Fig.3:**
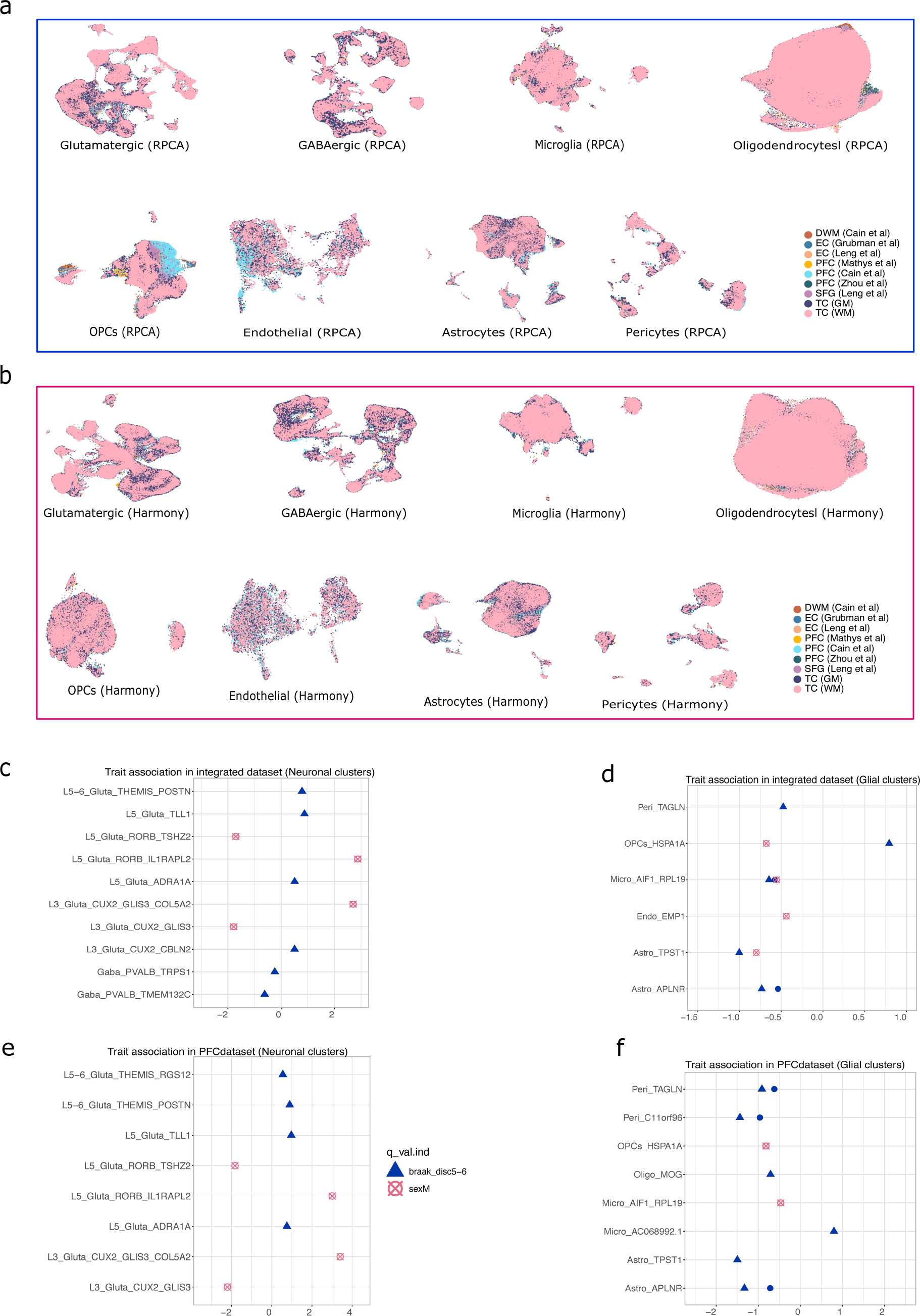
Integrating sub-cellular compositions and AD trait association analysis across multiple brain regions. **(a)** Integration of major cell types across multiple brain regions using RPCA(Seurat). **(b)** Integration of major cell types across multiple brain regions using Harmony. **(c)** Differential abundance of statistically significant neuronal subpopulations with respect to AD pathology (indicated by varying Braak stages) and sex across multiple brain regions. The x-axis represents the estimated log-ratio differences in relative abundance between different Braak stages and the y-axis represents different neuronal subpopulations. **(d)** Same as ‘c’ for glial subpopulations. **(e)** same as ‘c’ for neuronal subpopulations in PFC region only. **(f)** same as ‘c’ for glial subpopulations in PFC region only.

**Extended Data Fig.4:**
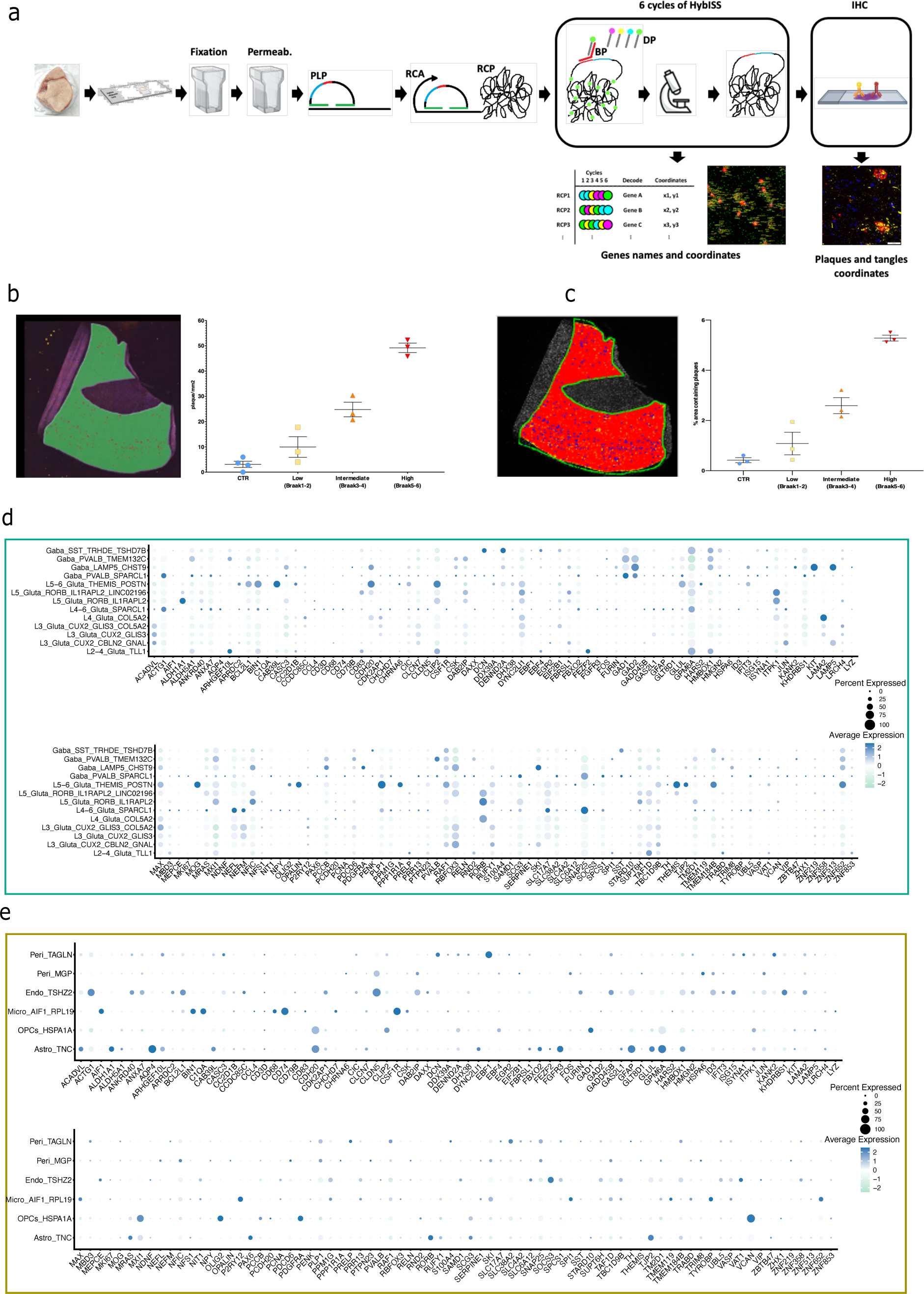
ISS workflow and 155 gene panel. **(a)** Workflow for combined *ISS* and *IHC.* Fresh frozen brain tissue blocks were sectioned, and sections were fixed with PFA and permeabilized using HCl before hybridization with custom made padlock probes (PLPs), followed by ligation, and rolling circle amplification (RCA) to generate rolling circle products (RCPs). Using a combination of bridge probes (BPs) and cycle-specific-fluorescent detection probes (DPs), 6 cycles of imaging were performed, followed every time by a round of stripping with formamide to remove the BP-DP complexes. The obtained images were used to generate spatially resolved expression profiles for the genes of interest. The BP-DP complexes were removed from the tissue sections, which were used to perform an immunohistochemistry to detect the localization of both β -amyloid plaques and tau tangles. (**b)** Detection of β -amyloid pathology (no. of b-amyloid plaques) using a classifier. **(c)** Classification of tiles based on presence/absence of plaques in defined areas (100×100 pixels). (**d-e),** Expression of 155 genes selected for CARTANA in neuronal and glial subpopulations respectively that were associated with AD pathology at single nucleus level.

**Extended Data Fig.5:**
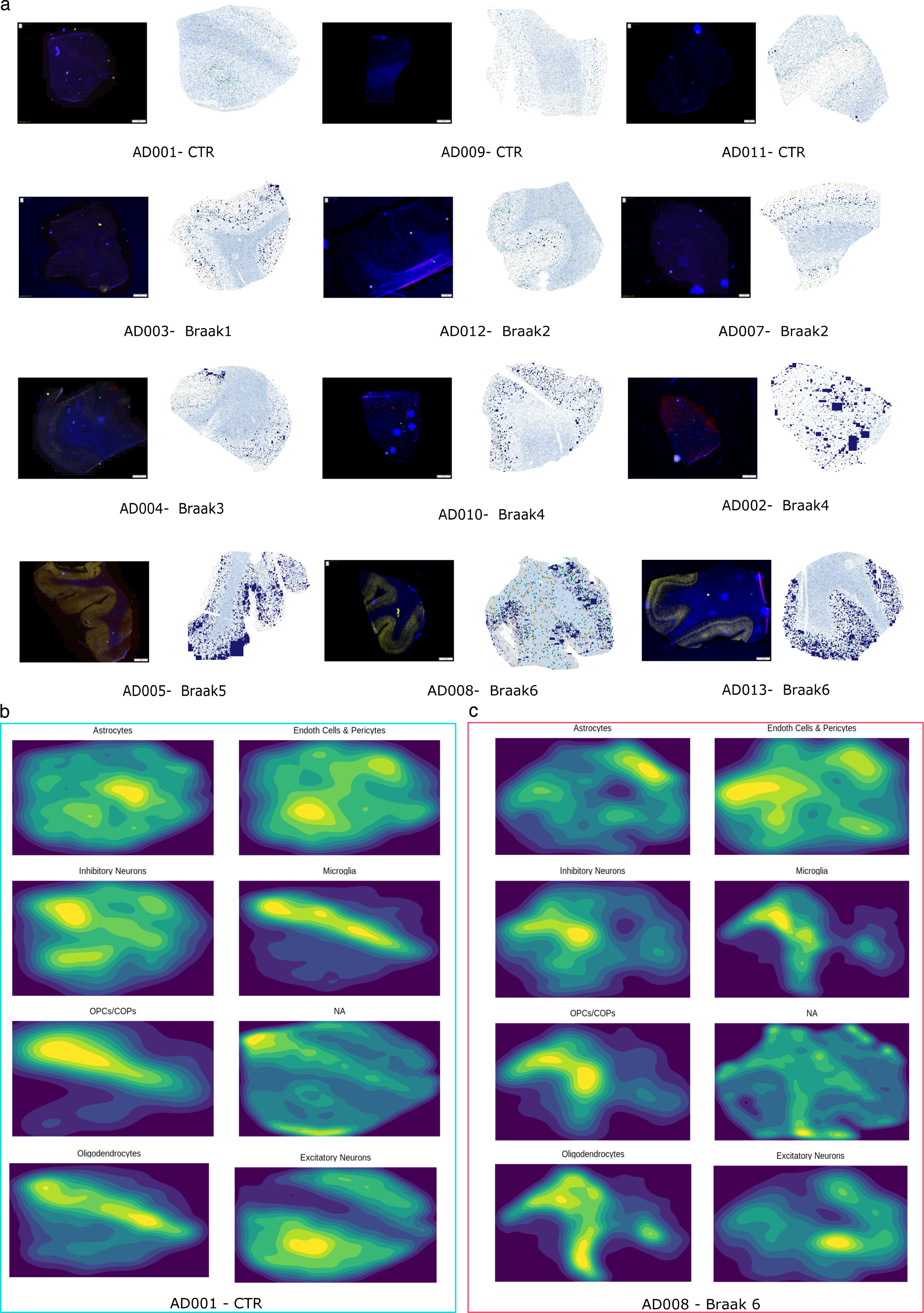
Immunohistochemistry of CARTANA samples. **(a)** IHC images of CARTANA samples and visual representation of the spatial distribution of cells same as Fig.4a-b. (**b-c)** Faceted 2D filled density plot where each facet corresponds to a different cell type. The filled areas in each plot represent regions of higher density in the spatial distribution of specific cell types. Panel ‘b’ represents a control sample while panel ‘c’ represents a sample with advanced AD pathology.

**Extended Data Fig.6:**
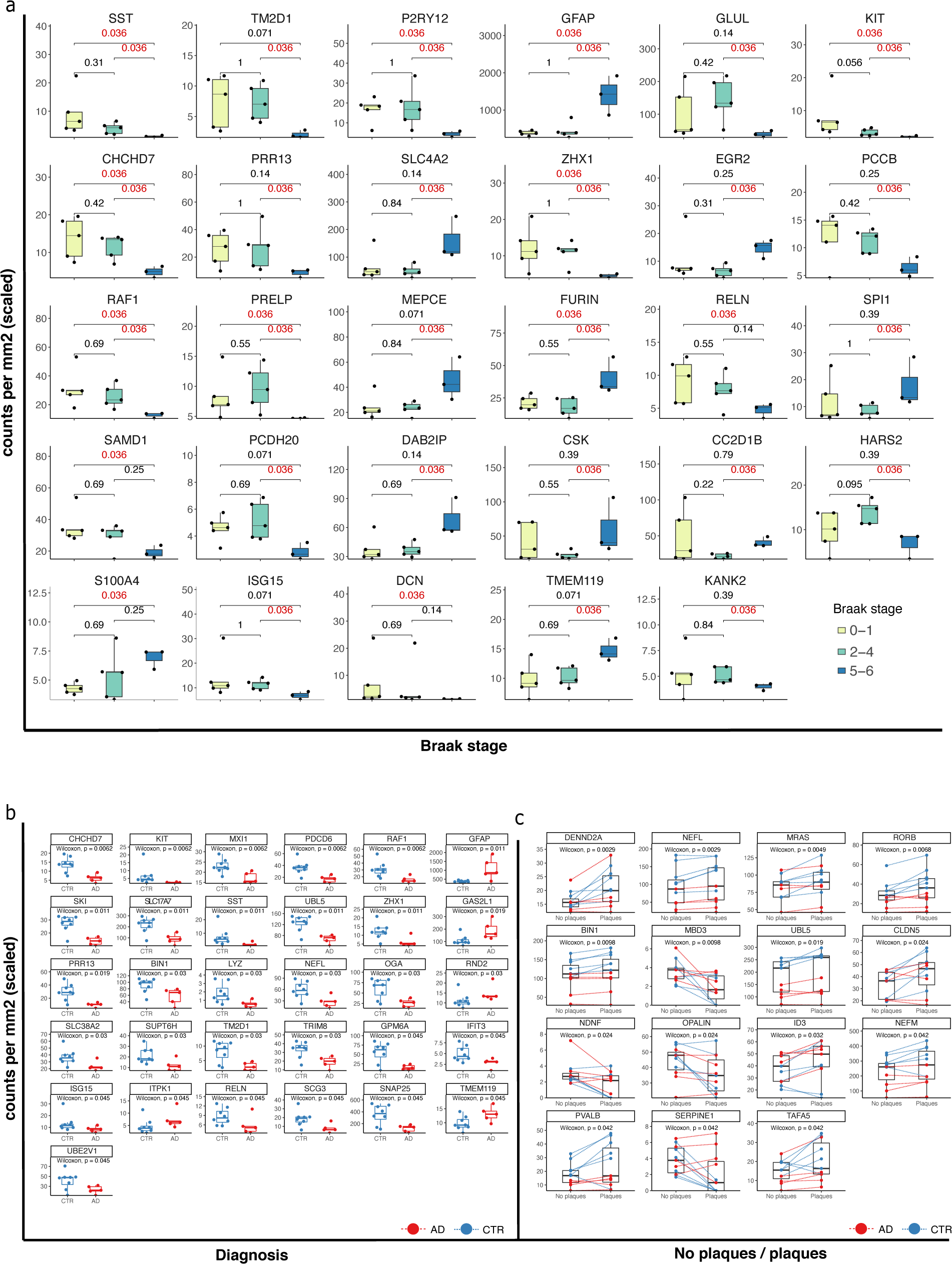
Comparisons of pathology indicators and controls. Boxplot of the number of normalized counts for selected genes (nominal p-value <0.05 in at least one comparison) at different **(a)** Braak stages, **(b)** Diagnosis (AD vs control), (**c)** No plaques vs plaques. All *p-values* were obtained using a Wilcoxon test for specified corrections and are not corrected for multiple testing.

**Extended Data Fig.7:**
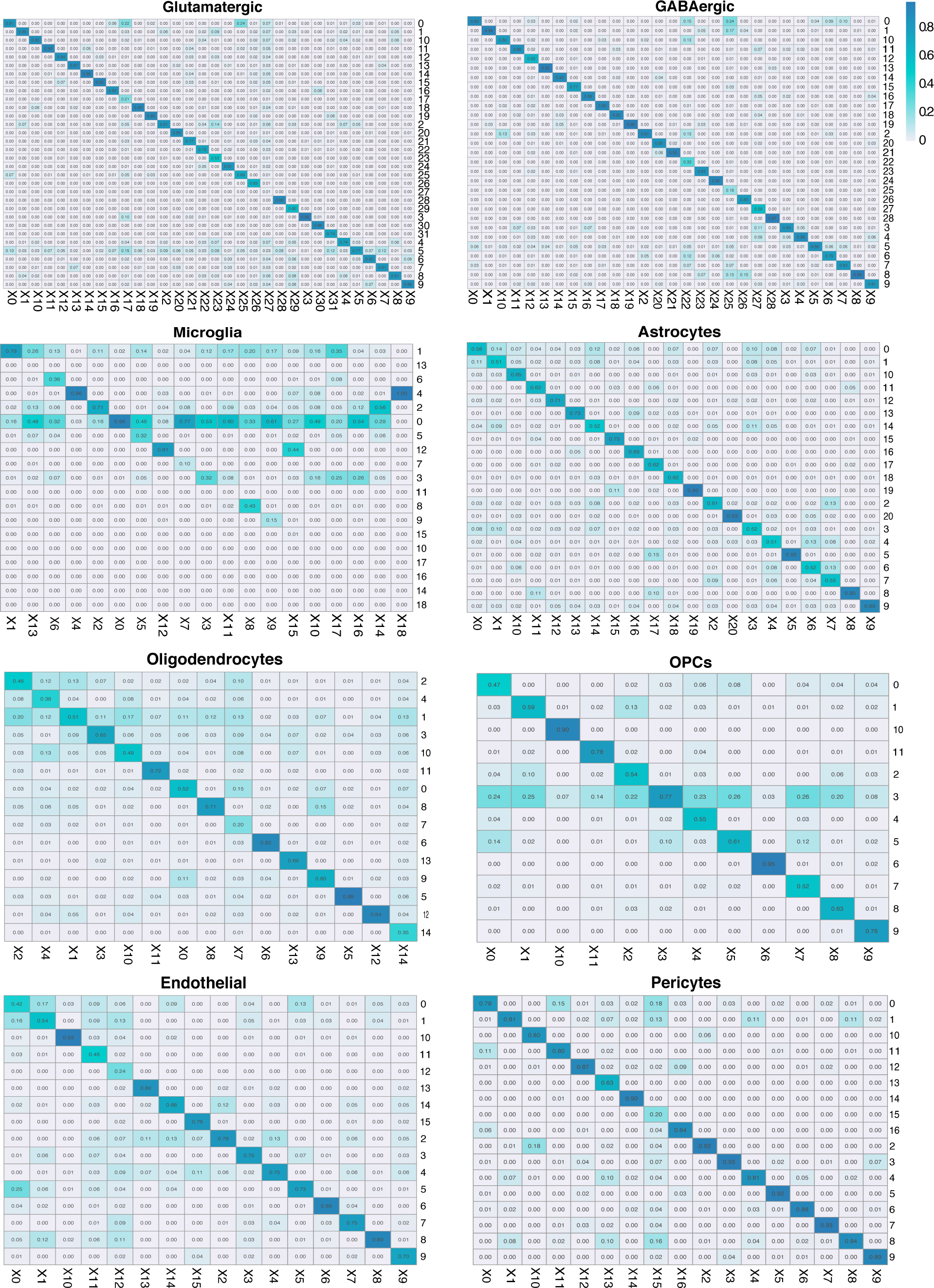
Machine learning for cluster stability. Confusion Matrix for random forest classifier for each cell type.

**Extended Data Fig.8:**
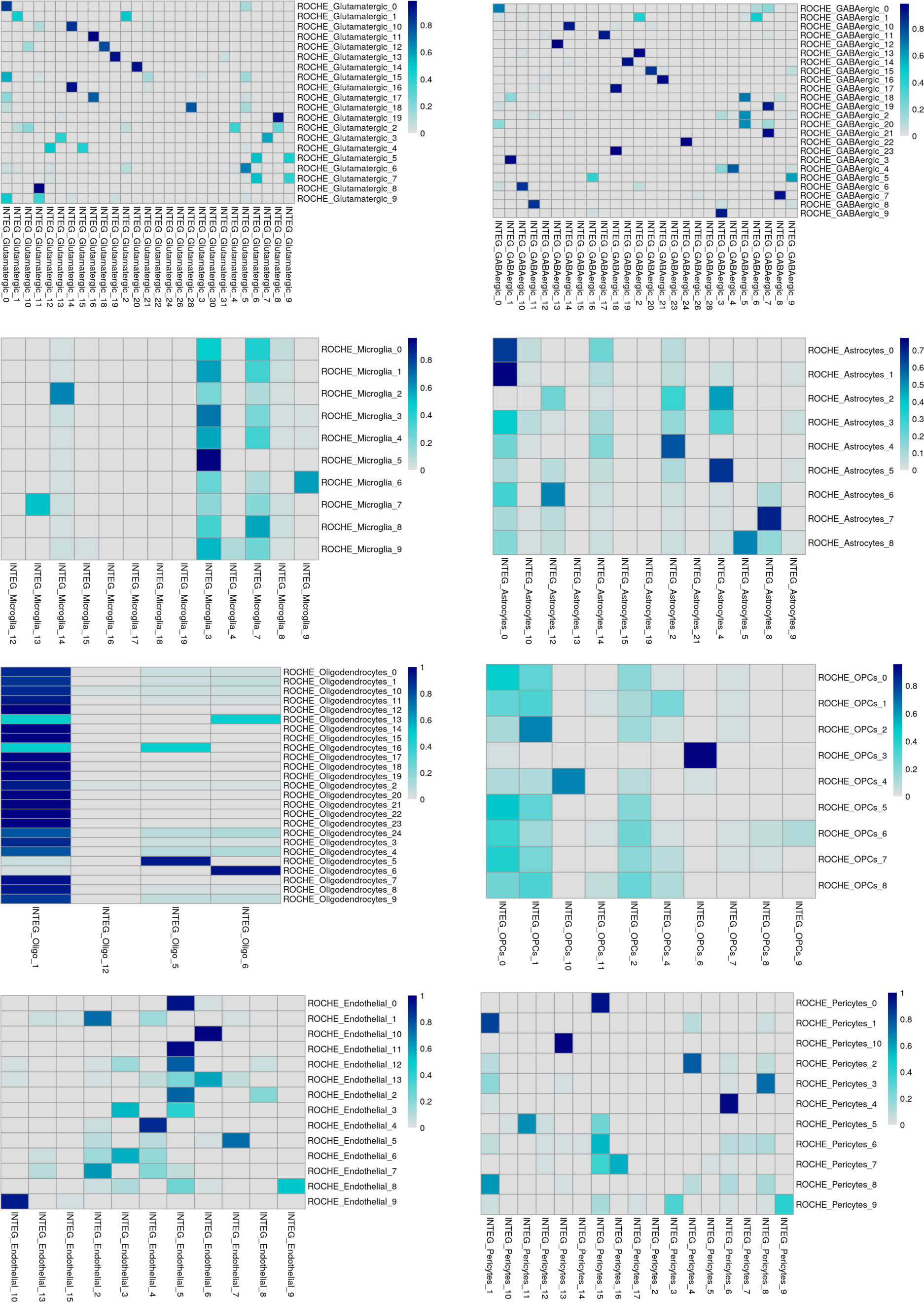
Cluster mapping of TC and integrated multiple brain regions. Comparison of TC region cluster and integrated multiple brain region for major cell types to finalize the cluster labels after merging similar subclusters.

## Acknowledgments

We extend our gratitude to the generous individuals who have donated their brains to further research at Netherlands Brain Bank. This work was supported in part by NIH grants R01AG066831 (VM) and U19AG074862 (VM).

## References

1. Mucke, L. Alzheimer’s disease. Nature 461, 895–897 (2009).

2. Dolgin, E. Alzheimer’s disease is getting easier to spot. Nature 559, S10–S12 (2018).

3. Mostafavi, S. et al. A molecular network of the aging human brain provides insights into the pathology and cognitive decline of Alzheimer’s disease. Nature Neuroscience 21, 811–819 (2018).

4. Williams, Cao & Yan. Transcriptomic analysis of human brains with Alzheimer’s disease reveals the altered expression of synaptic genes linked to cognitive deficits. Brain Communications 3, (2021).

5. Mathys, H. et al. Single-cell transcriptomic analysis of Alzheimer’s disease. Nature 570, 332–337 (2019).

6. Heneka, M. T. et al. Neuroinflammation in Alzheimer’s disease. The Lancet Neurology 14, 388–405 (2015).

7. Obulesu, M. & Jhansilakshmi, M. Neuroinflammation in Alzheimer’s disease: an understanding of physiology and pathology. International Journal of Neuroscience 124, 227–235 (2013).

8. Miyoshi, E. et al. Spatial and single-nucleus transcriptomic analysis of genetic and sporadic forms of Alzheimer’s Disease. 10.1101/2023.07.24.550282 (2023).

9. Luquez, T. et al. Cell type-specific changes identified by single-cell transcriptomics in Alzheimer’s disease. Genome Medicine 14, (2022).

10. Leng, K. et al. Molecular characterization of selectively vulnerable neurons in Alzheimer’s disease. Nature Neuroscience 24, 276–287 (2021).

11. Grubman, A. et al. A single-cell atlas of entorhinal cortex from individuals with Alzheimer’s disease reveals cell-type-specific gene expression regulation. Nature Neuroscience 22, 2087–2097 (2019).

12. Cain, A. et al. Multi-cellular communities are perturbed in the aging human brain and Alzheimer’s disease. Nature Neuroscience (2020) 10.1038/s41593-023-01356-x.

13. Zhou, Y. et al. Human and mouse single-nucleus transcriptomics reveal TREM2-dependent and TREM2-independent cellular responses in Alzheimer’s disease. Nature Medicine 26, 131–142 (2020).

14. Application Note: Mapping brain cell types with CARTANA in situ sequencing on the Nikon Ti2-E microscope. Nature https://www.nature.com/articles/d42473-019-00264-8.

15. Muratore, C. R. et al. Cell-type Dependent Alzheimer’s Disease Phenotypes: Probing the Biology of Selective Neuronal Vulnerability. Stem Cell Reports 9, 1868–1884 (2017).

16. Kong, W. et al. Independent component analysis of Alzheimer’s DNA microarray gene expression data. Molecular Neurodegeneration 4, (2009).

17. Squillario, M. & Barla, A. A computational procedure for functional characterization of potential marker genes from molecular data: Alzheimer’s as a case study. BMC Medical Genomics 4, (2011).

18. Cruchaga, C. et al. GWAS of Cerebrospinal Fluid Tau Levels Identifies Risk Variants for Alzheimer’s Disease. Neuron 78, 256–268 (2013).

19. Lai, M. K. P., Esiri, M. M. & Tan, M. G. K. Genome-wide profiling of alternative splicing in Alzheimer’s disease. Genomics Data 2, 290–292 (2014).

20. Jeong, Y. et al. 1950 MHz Electromagnetic Fields Ameliorate Aβ Pathology in Alzheimer’s Disease Mice. Current Alzheimer Research 12, 481–492 (2015).

21. Ali, F., Baringer, S. L., Neal, A., Choi, E. Y. & Kwan, A. C. Parvalbumin-Positive Neuron Loss and Amyloid-β Deposits in the Frontal Cortex of Alzheimer’s Disease-Related Mice. Journal of Alzheimer’s disease : JAD 72, 1323–1339 (2019).

22. Yin, G. N., Lee, H. W., Cho, J.-Y. & Suk, K. Neuronal pentraxin receptor in cerebrospinal fluid as a potential biomarker for neurodegenerative diseases. Brain Research 1265, 158–170 (2009).

23. Richens, J. L. et al. Practical detection of a definitive biomarker panel for Alzheimer’s disease; comparisons between matched plasma and cerebrospinal fluid. International journal of molecular epidemiology and genetics 5, 53–70 (2014).

24. Deng, Y. et al. Loss of LAMP5 interneurons drives neuronal network dysfunction in Alzheimer’s disease. Acta Neuropathologica 144, 637–650 (2022).

25. Cacabelos, R., Cacabelos, P. & Torrellas, C. Personalized Medicine of Alzheimer’s Disease. Handbook of Pharmacogenomics and Stratified Medicine 563–615 (2014) doi:10.1016/B978-0-12-386882-4.00027-X.

26. Libiger, O. et al. Identification of NPTX2 as a prognostic biomarker of Alzheimer’s disease through a longitudinal CSF proteomics study in ADNI subjects. Alzheimer’s & Dementia 16,.

27. Bryois, J. et al. Cell-type-specific cis-eQTLs in eight human brain cell types identify novel risk genes for psychiatric and neurological disorders. Nature Neuroscience 25, 1104–1112 (2022).

28. Yu, X., Ng, C. P., Habacher, H. & Roy, S. Foxj1 transcription factors are master regulators of the motile ciliogenic program. Nature Genetics 40, 1445–1453 (2008).

29. Faulty regulation of tau phosphorylation by the reelin signal transduction pathway is a potential mechanism of pathogenesis and therapeutic target in Alzheimer’s disease. European Neuropsychopharmacology 16, 547–551.

30. Steinberg, S. et al. Loss-of-function variants in ABCA7 confer risk of Alzheimer’s disease. Nature Genetics 47, 445–447 (2015).

31. Mathys, H. et al. Single-cell atlas reveals correlates of high cognitive function, dementia, and resilience to Alzheimer’s disease pathology. Cell 186, 4365–4385.e27 (2023).

32. Kocherhans, S. et al. Reduced Reelin Expression Accelerates Amyloid-Plaque Formation and Tau Pathology in Transgenic Alzheimer’s Disease Mice. Journal of Neuroscience 30, 9228–9240 (2010).

33. Horiuchi, K. et al. Identification and Characterization of a Novel Protein, Periostin, with Restricted Expression to Periosteum and Periodontal Ligament and Increased Expression by Transforming Growth Factor β. Journal of Bone and Mineral Research 14, 1239–1249 (1999).

34. Shih, C.-H., Lacagnina, M., Leuer-Bisciotti, K. & Pröschel, C. Astroglial-Derived Periostin Promotes Axonal Regeneration after Spinal Cord Injury. The Journal of Neuroscience 34, 2438–2443 (2014).

35. Shimamura, M. et al. Role of Central Nervous System Periostin in Cerebral Ischemia. Stroke 43, 1108–1114 (2012).

36. Matsunaga, E. et al. Periostin, a neurite outgrowth-promoting factor, is expressed at high levels in the primate cerebral cortex. Development, Growth & Differentiation 57, 200–208 (2015).

37. Yu, Y. et al. Comprehensive RNA-Seq transcriptomic profiling across 11 organs, 4 ages, and 2 sexes of Fischer 344 rats. Scientific Data 1, (2014).

38. De Jager, P. L. et al. A genome-wide scan for common variants affecting the rate of age-related cognitive decline. Neurobiology of Aging 33, 1017.e1–1017.e15 (2012).

39. Laywell, E. D. et al. Enhanced expression of the developmentally regulated extracellular matrix molecule tenascin following adult brain injury. Proceedings of the National Academy of Sciences 89, 2634–2638 (1992).

40. Green, G. S. et al. Cellular dynamics across aged human brains uncover a multicellular cascade leading to Alzheimer’s disease. 10.1101/2023.03.07.531493 (2023).

41. Germain, P.-L., Lun, A., Meixide, C. G., Macnair, W. & Robinson, M. D. Doublet identification in single-cell sequencing data using scDblFinder. F1000Research 10, (2022).

42. Barkas, N. et al. Joint analysis of heterogeneous single-cell RNA-seq dataset collections. Nature Methods 16, 695–698 (2019).

43. Korsunsky, I. et al. Fast, sensitive and accurate integration of single-cell data with Harmony. Nature Methods 16, 1289–1296 (2019).

44. Lun, A. T. L., McCarthy, D. J. & Marioni, J. C. A step-by-step workflow for low-level analysis of single-cell RNA-seq data with Bioconductor. F1000Research 5, 2122 (2016).

45. Amezquita, R. A. et al. Orchestrating single-cell analysis with Bioconductor. Nature Methods 17, 137–145 (2019).

46. Stuart, T. et al. Comprehensive Integration of Single-Cell Data. Cell 177, 1888–1902.e21 (2019).

47. dittoSeq. Bioconductor https://bioconductor.org/packages/dittoSeq.

48. Lin, H. & Peddada, S. D. Analysis of compositions of microbiomes with bias correction. Nature Communications 11, (2020).

49. Gelman, A. Scaling regression inputs by dividing by two standard deviations. Statistics in Medicine 27, 2865– 2873 (2007).

50. Robinson, M. D., McCarthy, D. J. & Smyth, G. K. edgeR: a Bioconductor package for differential expression analysis of digital gene expression data. Bioinformatics 26, 139–140 (2009).

51. Brooks, M., E., et al. glmmTMB Balances Speed and Flexibility Among Packages for Zero-inflated Generalized Linear Mixed Modeling. The R Journal 9, 378 (2017).

52. Robinson, M. D. & Oshlack, A. A scaling normalization method for differential expression analysis of RNA-seq data. Genome Biology 11, R25 (2010).

53. de Leeuw, C. A., Mooij, J. M., Heskes, T. & Posthuma, D. MAGMA: Generalized Gene-Set Analysis of GWAS Data. PLOS Computational Biology 11, (2015).

54. Wightman, D. P. et al. A genome-wide association study with 1,126,563 individuals identifies new risk loci for Alzheimer’s disease. Nature Genetics 53, 1276–1282 (2021).

55. Korotkevich, G., et al. Fast gene set enrichment analysis. 10.1101/060012 (2016).

